# Process engineering of biopharmaceutical production in moss bioreactors via model-based description and evaluation of phytohormone impact

**DOI:** 10.1101/2021.12.19.473340

**Authors:** Natalia Ruiz-Molina, Juliana Parsons, Sina Schroeder, Clemens Posten, Ralf Reski, Eva L. Decker

## Abstract

The moss Physcomitrella is an interesting production host for recombinant biopharmaceuticals. Here we produced MFHR1, a synthetic complement regulator which has been proposed for the treatment of diseases associated to the complement system as part of human innate immunity. We studied the impact of different operation modes for the production process in 5 L stirred-tank photobioreactors. The total amount of recombinant protein was doubled by using fed-batch or batch compared to semi-continuous operation, although the maximum specific productivity (mg MFHR1/g FW) increased just by 35%. We proposed an unstructured kinetic model which fits accurately with the experimental data in batch and semi-continuous operation under autotrophic conditions with 2% CO_2_ enrichment. The model is able to predict recombinant protein production, nitrate uptake and biomass growth, which is useful for process control and optimization. We investigated strategies to further increase MFHR1 production. While mixotrophic and heterotrophic conditions decreased the MFHR1-specific productivity compared to autotrophic conditions, addition of the phytohormone auxin (NAA, 10 µM) to the medium enhanced it by 470% in shaken flasks and up to 230% and 260%, in batch and fed-batch bioreactors, respectively. Supporting this finding, the auxin-synthesis inhibitor L-Kynurenine (100 µM) decreased MFHR1 production significantly by 110% and 580% at day 7 and 18, respectively. Expression analysis revealed that the MFHR1 transgene, driven by the Physcomitrella *actin5* (*PpAct5*) promoter, was upregulated 16 hours after NAA addition and remained enhanced over the whole process, whereas the auxin-responsive gene *PpIAA1A* was upregulated within the first two hours, indicating that the effect of auxin on *PpAct5* promoter-driven expression is indirect. Furthermore, the day of NAA supplementation was crucial, leading to an up to 8-fold increase of MFHR1-specific productivity (0.82 mg MFHR1/ g fresh weight, 150 mg accumulated over 7 days) compared to the productivity reported previously. Our findings are likely to be applicable to other plant-based expression systems to increase biopharmaceutical production and yields.

## 1 Introduction

The production of biopharmaceuticals, which include enzymes, vaccines, antibodies, growth factors, and hormones, is a complex task where small differences in production conditions can influence product quality, efficacy, safety and yield. As most commercial biopharmaceutical proteins are complex glycoproteins which cannot be produced in microorganisms, more than 50% are produced in mammalian cells (Tripathi and Shrivastava, 2019). Plant-based biopharmaceuticals have gained increasing attention as an alternative to mammalian cell systems in the last decade; however, only β-glucocerebrosidase to treat Gaucher disease (Elelyso®) produced in carrot cell suspensions is currently available on the market (van Dussen et al., 2013).

One advantage of plant-based expression systems is biosafety due to lack of contamination with animal-borne viruses. Furthermore, there are technical advantages associated with up-stream processing: For instance, a rapid scale-up of production using *Nicotiana benthamiana* and a transient expression system is feasible within two months (Shoji et al., 2012; Capell et al., 2020), which would allow a quick response to address a public health crisis. During the current COVID-19 pandemic, it became evident that diagnostic reagents and vaccine production capacity was not sufficient to meet demand, and transient expression in plants could compensate this (Tusé et al., 2020). The plant-derived Virus-Like Particles vaccine candidate for COVID-19 by Medicago completed phase I clinical trials with promising results and the phase II/III trials are ongoing (Gobeil et al., 2021; Ward et al., 2021). There are many plant-based vaccines in phase I clinical trials (Shim et al., 2019), and some of them completed phases II and III such as avian (monovalent) and seasonal (quadrivalent) influenza vaccines produced in *N. benthamiana* (Ward et al., 2020).

Plant cell-suspension cultures in bioreactors have advantages over whole-plant systems, because biopharmaceuticals can be produced under controlled and reproducible conditions, while complying with Good Manufacturing Practice (GMP) requirements. Furthermore, down-stream processing (DSP) may be simplified, thus reducing production time and costs (Huang and McDonald, 2009). Among other features, the high rate of homologous recombination observed in the nuclear DNA of somatic cells positioned the moss *Physcomitrium patens* (Physcomitrella) as an attractive platform for the production of recombinant therapeutic proteins (Decker et al., 2014; Reski et al., 2018; Bohlender et al., 2020).

Physcomitrella can be cultivated in suspensions in a differentiated, genetically stable, filamentous stage, the protonema tissue. This haploid tissue consists of two cell types: chloronema and caulonema. Chloronemal cells are characterized by cross-walls perpendicular to the growth axis of the filament and are rich in chloroplasts, while caulonemal cells are narrower with oblique cross-walls and fewer chloroplasts (Reski, 1998). Both cell types expand by tip growth with different rates, chloronemal cells divide every 24 h, caulonemal cells every 7 h (Schween et al., 2003). The transition from chloronema to caulonema can be triggered among others by glucose or the phytohormone auxin (Thelander et al., 2005). Glucose is coupled to developmental progression of Physcomitrella via cyclin D (Lorenz et al., 2003), and auxin is a signaling molecule involved in many developmental processes in plants including mosses (Decker et al., 2006).

Protonemal tissue grows under autotrophic conditions in inorganic media. Furthermore, the bioprocess has been scaled up to 500 L in wave bag photo-bioreactors and stirred tank photo-bioreactors complying with GMP conditions (Reski et al., 2015; Decker and Reski, 2020). The first recombinant pharmaceutical protein produced in moss which completed clinical trial phase I is α-galactosidase (Repleva aGal; eleva GmbH) to treat Fabry disease (Hennermann et al., 2019). In addition, α-glucosidase to treat Pompe disease (Repleva GAA, eleva GmbH) completed preclinical studies (Hintze et al., 2020). Both products showed superior characteristics compared to the mammalian cell-based products and are catalogued as biobetters (Hennermann et al., 2019; Hintze et al., 2020).

One of the bottlenecks of plant-based production is the low yield of the recombinant protein (Schillberg and Finnern, 2021). Albeit the cell-specific protein productivity of plant cells is comparable to CHO cells, the larger size of plant cells results in much lower volumetric productivity (Havenith et al., 2014). Protein yields generally range from 0.01-200 mg/L (Xu et al., 2011). For example, antibodies produced in tobacco leaves by transient expression yielded up to 2 mg/g FW (Zischewski et al., 2016) and a peptide vaccine produced in chloroplasts from tobacco yielded up to 7 mg/g FW (Molina et al., 2004). However, most complex biopharmaceuticals cannot be targeted to the chloroplasts but should be targeted to the secretory pathway, where they undergo posttranslational modifications. Therefore, many products could not yield more than 100 µg/g FW yet (Schillberg et al., 2019; Schillberg and Finnern, 2021). Many efforts have been undertaken to increase the yield in different stages of the bioprocess, such as optimization of the gene sequence, search for suitable promoters and terminators, design of synthetic 5’and 3’unstranslated regions (UTRs), insertion of multiple copies of the transgene into the genome, targeting strategies, co-expression with protease inhibitors, or suppression of protease gene expression, among others (Rozov and Deineko, 2019). Culture conditions also influence protein stability and productivity, which can be enhanced by optimizing physical parameters such as light, pH, temperature, agitation speed, aeration or nutritional requirements, bioreactor design and operating mode (batch, fed-batch, semi-continuous, continuous, perfusion). For instance, protein productivity increased between 1.2 and 25-fold in plant cell suspension cultures by changing bioreactor operating modes (Huang et al., 2010; Huang and McDonald, 2012).

Most approaches undertaken to increase recombinant protein yield in Physcomitrella have been focused on molecular strategies (Horstmann et al., 2004; Baur et al., 2005; Jost et al., 2005; Saidi et al., 2005; Schaaf et al., 2005; Weise et al., 2006; Peramuna et al., 2018; Top et al., 2021), while bioreactor engineering approaches and culture conditions have received less attention. Although moss growth and differentiation have been studied in different bioreactors and conditions (Hohe et al., 2002; Hohe and Reski, 2005; Lucumi et al., 2005; Lucumi and Posten, 2006; Cerff and Posten, 2012a, 2012b; Ruiz-Molina et al., 2016; Heck et al., 2021), the effects of carbon source (autotrophic, mixotrophic, heterotrophic), environmental conditions or bioreactor operation mode on recombinant protein yield are not reported in the literature.

The need for optimization and better process control in the biopharmaceutical industry has increased during the last years due to the rise in demand of these drugs and more competitive markets (López-Meza et al., 2016). Mathematical models in biotechnology are a powerful tool for rational and straight-forward process development thus saving time and resources. Among these are phenomenological models which derived mainly from microbial systems, and need understanding of the biochemical process and relationship between macroscopic variables such as biomass, substrate and product (Alvarez, 2014). These models are also widely applied to optimize processes for biofuels and high-value compounds produced by microalgae, such as carotenoids, phycobilins and fatty acids, e.g. eicosapentenoic acid and docosahexaenoic acid (DHA) (Lee et al., 2015). Dynamic models to predict and optimize recombinant protein production are less exploited in molecular farming (biopharmaceutical production), probably because transient platforms often lack reproducibility between batches, due to high variability in expression levels (Alvarez, 2014). A model based on a design-of-experiments approach was implemented to solve the uncertainty and predict the amount of recombinant protein produced in these platforms by identifying the main factors involved in recombinant protein productivity (Buyel and Fischer, 2012). Plant cell-suspension cultures can also benefit from mathematical modelling to optimize the process with less experimental effort, however, just a few studies have applied them (Alvarez, 2014). Nevertheless, so far there are no dynamic models for recombinant protein production in the moss bioreactor.

Here, we aimed at increasing the recombinant protein yield in moss bioreactors. As a case study, we produced the complex glycoprotein MFHR1, a synthetic complement regulator (Michelfelder et al., 2018), in Physcomitrella. The human complement system constitutes a crucial part of innate immunity and protects the body from invading pathogens (Merle et al., 2015). MFHR1 is a fusion protein, which combines the regulatory activity of human complement factor H (FH) and FH-related protein 1. Alongside moss-produced FH, it is considered a potential therapeutic agent against complement-associated diseases such as C3 glomerulopathies (C3G), atypical hemolytic uremic syndrome (aHUS), or viral infections where complement over-activation has been associated to the pathogenesis, such as COVID-19 or severe dengue fever (Michelfelder et al., 2017, 2018; Top et al., 2019; Ruiz-Molina et al., 2021). In our present study, we produced two MFHR1 variants, differing in one amino acid at position 62 of FH equivalent to position 193 of MFHR1 (V62I) (Ruiz-Molina et al., 2021).

Due to a high variation in MFHR1 yields between transgenic moss lines, we evaluated if product yield positively correlates with mRNA expression levels and transgene copies integrated into the genome. Our findings suggest that multiple transgene copies are desirable to maximize protein yields in Physcomitrella. We explored the effect of process operating mode (batch, fed-batch, and semi-continuous) on MFHR1 production as a means for targeted changes of environmental conditions for the cells. Based on these experiments we propose a kinetic model of the moss bioreactor which can accurately predict recombinant protein production, biomass growth and nitrate uptake under batch and semi-continuous operation, and can be used to maximize protein and biomass productivity. This model contributes to a deeper understanding of a plant-based system and dynamic changes of the variables. Moreover, we explored the effect of the crucial plant growth regulator auxin on recombinant protein yield and its influence on actin gene expression. The kinetic model and the positive influence of auxin on biopharmaceutical production might be applied to other plant-based systems aiming at increasing biopharmaceutical yields.

## 2. Materials and Methods

### 2.1 Protonema suspension cultures

Previously established transgenic moss lines P1 and N-179 producing MFHR1^V62^ and MFHR1^I62^ variants, respectively, were used (Top et al., 2019; Ruiz-Molina et al., 2021). These variants are characterized by valine or isoleucine in position 193, corresponding to position 62 of FH, which lead to a difference in their activity (Ruiz-Molina et al., 2021). For simplification and better understanding of the results, P1 is called line A and N-179 line B.

Suspension cultures were initiated with a cell density around 100 mg dry weight (DW)/ L in 100 mL shaken flasks (25 mL working volume). 2-(N-morpholino)ethanesulfonic acid (MES) was added and the pH was set to 5.5. To determine the correlation between copy number of integrated expression cassettes, mRNA expression and protein production, transgenic moss lines were propagated in suspension cultures during one year. Cell suspension cultures were initiated with around 100 mg dry weight (DW)/ L and harvested at day 18.

### 2.2 Description of the mathematical model

The model developed is built up from a physiological level represented by formal kinetics for the specific growth rates. Material balances are employed on the macroscopic level to describe the dynamics of biomass (c_x_), nitrate (c_N_) and recombinant protein (c_p_) concentrations. r_X_, r_N_, r_p_ define the specific growth rate, the specific substrate uptake rate and the specific recombinant protein production rate, respectively [d^-1^].

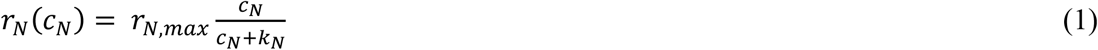

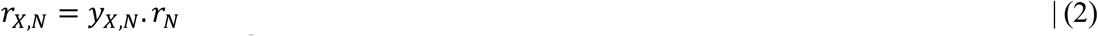

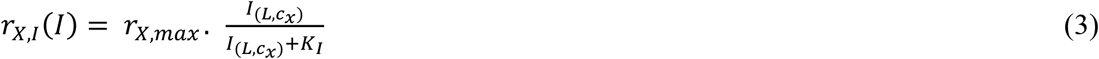

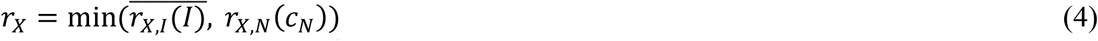

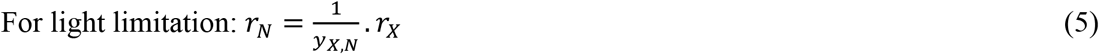

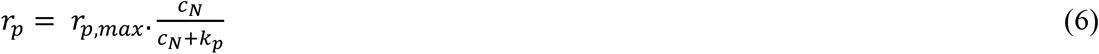

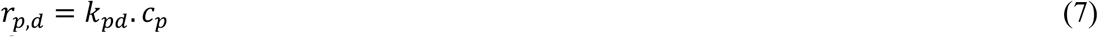

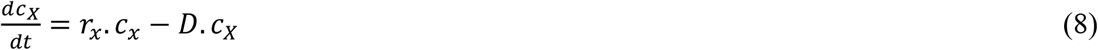

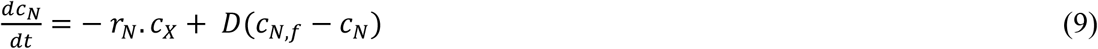

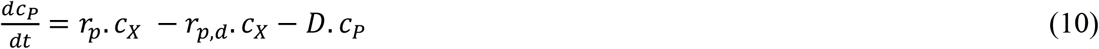

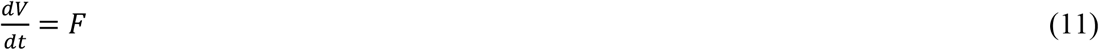

Equations 1-5 describe the specific growth and substrate uptake rates. One important factor in autotrophic organisms is the light. Limitation of nutrients and light have been widely reported in microalgae (Béchet et al., 2013; Lee et al., 2015). To construct this kinetic model, just nitrate and light were taken into account since these are the first to be limited in Physcomitrella cultures under our conditions. This co-limitation was described using the so-called minimum law or threshold model, in which the growth is affected by the most limited substrate (eq. 4) (Lee et al., 2015). The most common models for growth and substrate consumption correspond to Monod type (Monod, 1949), which introduced the concept of a growth-limiting substrate. Further development has been introduced by Jhon Pirt interpreting substrate uptake as enzymatic step combined with Michaelis-Menten kinetics and in a second step a yield with which the cell can support growth based on the substrate taken up. In this way also an equation for the substrate uptake rate (r_N_) is delivered (eq 1-2 for nitrate limitation and eq 3-5 for light limitation), where y_x,N_ is the biomass yield coefficient (g biomass/ g nitrate consumed), and k_i_, and k_N_ are the saturation constants of the limiting substrates, light and nitrate, respectively [g/L]. Nitrate used for non-growth associated purposes (maintenance term) was neglected. Monod-like equations have been used to model growth kinetics in autotrophic processes, where light, the energy source, is considered a substrate (Béchet et al., 2013). The growth kinetics of Physcomitrella under different light intensities suggested Han Model behavior (Cerff and Posten, 2012b; Schediwy et al., 2019); therefore, we incorporated Monod’s equation in the threshold model to explain the substrate consumption and specific growth rate (eq 3).

Modelling of microalgae processes generally uses the Beer-Lambert law to predict the light distribution in the bioreactor (Béchet et al., 2013). This equation describes an exponential decay of light intensity from the external surface, which is partially compensated by the geometric intensification against the axis of the cylinder. This results in a reasonable constant light intensity in the reactor. Beer-Lambert assumes that the cells do not scatter the light, which does not reflect reality. Although more complex models exist, they are difficult to implement and are computationally intensive (Dauchet et al., 2013), however, the Beer-Lambert law could be accurate enough to describe light distribution in the bioreactor (Luo and Al-Dahhan, 2012). Therefore, we implemented this equation in our model, assuming that the reactor surface is homogenously illuminated, following the methodology described by Evers (1991) (Supplementary eq 1-5), where l_0_ is the incident light [µmol/(m^2^s)], σ_x_ is the cell absorption coefficient [m^2^/g] and r_R_ is the cylinder radius [m]. The path length of light (p) is function of a distance from vessel surface (L) and angle of light path (Θ) (Supplementary eq 2). I(L,c_X_) is the mean light intensity as a function of biomass concentration (c_x_) and location in the vessel. It takes into account light flux from all directions (from 0 to *π* due to symmetry) (Supplementary eq 3). The overall biomass growth rate considering the light as the limiting substrate 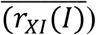 is calculated with supplementary equation 4, combining supplementary equations 3 and 2, where r_X,max_ is the maximum specific growth rate [d^-1^] (Supplementary eq 5).

The nitrogen source is essential for protein production, therefore we used equation 5 to simulate recombinant protein production originated from Monod model. r_p,max_ represents the maximum specific recombinant protein production rate [d^-1^], k_p_ is nitrate saturation constant for protein of interest synthesis [g/L]. The degradation rate (r_p,d_) [d^-1^] was also taken into account, depending on biomass and product concentration (eq 6).

The equations 7-10 describe a batch, fed-batch or semi-continuous process, where D is the dilution rate, and dV/dt is the dynamics of volume in the bioreactor. D is zero in a batch run and dV/dt is zero in semi-continuous and batch operations. c_N,f_ is the substrate concentration in the feeding stream in a fed-batch or semi-continuous operation.

### 2.3 Parameter estimation

Ordinary differential equations (ODE) were solved by the numeric method Runge-Kutta, using the solver solve_IVP implemented in SciPy (Python) (Virtanen et al., 2020). Estimation of parameters was performed by minimizing the distance between experimental data (Y_ij_) and the ODE solution (*function*(*X*_*i*_,*P*_*k*_)). The minimization of the objective or error function (least square minimum) (eq 12) was performed by using differential evolution, a global optimization method implemented in SciPy (Python).

### 2.4 Bioreactor cultures

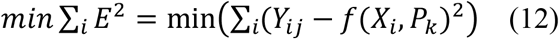

The cultures were scaled up to stirred tank photobioreactor (5L) with Knop ME medium using the following conditions: aeration: 0.3 vvm (2% CO_2_), agitation with pitched 3 blade impeller at 500 rpm under continuous light. The light was increased from 160 to 350 µmol/m^2^ s after 2 days. The illumination using LEDs were evenly distributed on the surface of the bioreactor, the pH was controlled at 4.5 and temperature was kept at 22°C.

For fed-batch operation the process was carried out in batch with 4L volume for 7 days and the feeding was started at day 7. The feeding flow (F) with 5x concentrated Knop ME medium was set to 200 mL/d.

For the semi-continuous operation, the process was carried out in batch with 5 L volume for 5 or 6 days, respectively, and the semi-continuous phase started at day 6 or 7. For this, 2 L or 1 L suspension were harvested and replaced with fresh Knop ME medium daily, which correspond to a dilution rate of 0.4 or 0.2 d^-1^, respectively.

### 2.5 Auxin supplementation and auxin biosynthesis inhibition

For the supplementation with the auxin 1-Naphtalene acetic acid (NAA, Sigma), 100 mM stock solution was prepared in 0.5M KOH. The auxin biosynthesis inhibitor, L-kynurenine, (Sigma) was prepared in DMSO.

### 2.6 Determination of biomass dry weight, nitrate concentration and protein accumulation

For dry weight (DW) measurement, 10-50 mL of tissue suspension were vacuum-filtered and dried for 2 h at 105 °C. For experiments at shaken-flask scale, growth index was calculated as (initial biomass-final biomass)/ initial biomass. Nitrate concentration in the culture supernatant was measured in the reflectometer (RQflex®) using the Reflectoquant® Nitrate Test following the manufacturer’s instructions. During bioreactor runs, nitrate was also measured at day 0, because it differs slightly between batches, due to experimental error.

MFHR1 production was analyzed from the cellular fraction (specific productivity µg MFHR1/ g FW). The protein was extracted from 30 mg moss fresh weight (FW) (vacuum-filtered material and frozen in liquid nitrogen). The tissue was homogenized and disrupted using two beads (glass and metal) in a tissue-lyzer for 1 min at 30 Hz, followed by addition of the extraction buffer (4 µL /mg FW) (60 mM Na_2_HPO_4_ x 2 H_2_O, 60 mM KH_2_PO_4_, 408 mM NaCl, 60 mM EDTA, 1% protease inhibitor (P9599, Sigma-Aldrich)) and by ultrasonic bath for 10 min. Protein was quantified in the extraction supernatant by ELISA as described previously (Ruiz-Molina et al 2021). Briefly, microtiter plates (Nunc Maxisorp, Thermo Fisher Scientific) were coated overnight with a monoclonal antibody recognizing the SCR 20 of FH (GAU 018-03-02, Thermo Fisher Scientific), diluted 1:2,000 in coating buffer (1.59 g/l Na_2_CO_3_, 2.93 g/l NaHCO_3_, pH 9.6). The protein was detected using a polyclonal anti-SCR 1-4 (1:15,000 in Tris Buffer Saline (TBS) supplemented with 1% BSA and 0.05% Tween-20) (Kühn et al., 1995) and anti-rabbit horseradish peroxidase (HRP) (NA934; Cytiva) (diluted 1:5,000 in washing buffer).

### 2.7 Nucleic acids extraction and quantitative PCR

Auxin treated samples were harvested from the bioreactor. The suspension was vacuum-filtered and 30 mg FW plant material frozen in liquid nitrogen. The frozen tissue was homogenized and disrupted using two beads (glass and metal) in a tissue-lyzer for 1 min at 30 Hz. RNA was extracted using RNeasy plant mini kit (Qiagen). After DNAse I treatment, cDNA synthesis was carried out from 1 µg RNA using Multiscribe RT and TaqMan Reverse Transcription reagents (Thermo Fisher Scientific) and a control of complete DNA digestion was included without addition of Multiscribe RT. Quantitative RT-PCR was performed with 10 ng/µl cDNA, 400 nM of each primer and 1x SensiFast SYBR® No-ROX (Bioline) in the LightCycler 480 System (Roche) under the following conditions: 95°C for 2 min (one cycle); 95°C for 5 s, 60°C for 10 s, and 72°C for 15 s (45 cycles). A thermal ramping stage was included to obtain melting curves.

Primers were designed using the Universal Probe Library Assay Design Center (Roche) and checked for specificity against the Physcomitrella transcriptome in the Phytozome database (Goodstein et al., 2012) release v13. Every primer pair was tested for specificity and efficiency using serial dilutions DNA. Primers with an efficiency lower than 1.9 were discarded. Expression levels of the transgene (*MFHR1*), *PpIAA1A* (Pp3c8_14720V3.1), and actin 7 *(PpAct7a)* (Pp3c3_33410V3.1), were analyzed using the primers listed in Supplementary Table S1.

Normalized expression was calculated according to the ΔΔC_T_ method, using the Light Cycler 480 software (Roche). Genes coding for elongation factor 1-α (EF1-α) (Pp3c2_10310V3.1) and 60S ribosomal protein L21 (Pp3c13_2360.V3.1) were used as housekeeping references to normalize the data. Relative quantification was performed using a time point before addition of NAA in the bioreactor as calibrator. Every sample was tested in triplicates.

The number of integrations of the *MFHR1* transgene in the genome was estimated by qPCR as described above. DNA was isolated using GeneJet Plant Genomic DNA Purification Mini Kit (Thermo Scientific) according to the manufacturer’s instructions. The single copy gene *PpCLF* (Pp3c22_22940V3.1) was used as a reference to normalize the data using two primer pairs listed in Supplementary Table S1. The number of transgene copies was determined using 2 controls as calibrators, a moss line with a single integration of the hygromycin resistance gene (*hpt*), and the parental line (*Δxt/ft*) which has a single integration of the 5’HR region included in the construct (Ruiz-Molina et al., 2021). The *MFHR1* transgene was amplified with the same primers used for the qRT-PCR. All primers are listed in Supplementary Table S1.

### 2.8 Statistics

Analyses and figures were performed with the GraphPad Prism software version 8.0 for Windows (GraphPad software, San Diego, California, USA). Statistical significance was evaluated by ANOVA followed by Bonferroni or Tukey test (p<0.05).

## 3 Results and Discussion

### 3.1 A high copy number of integrated expression cassettes is important for high MFHR1 productivity

MFHR1 is considered a potential therapeutic agent for complement-associated diseases. We produced two variants of this synthetic complement regulator, characterized by an amino acid exchange (V193I), derived from the FH haplotype (V62I) (Ruiz-Molina et al., 2021).

First, we studied the correlation between copy number of integrated expression cassettes, mRNA expression and protein production for 14 MFHR1-producing moss lines. We analyzed the transgenic plants after a long period of cultivation (1 year after transfection) to avoid episomal expression (Murén et al., 2009). DNA-copy number, mRNA levels, and recombinant protein yields were analyzed by qPCR, qRT-PCR and ELISA, respectively. It is important to mention that under our conditions, MFHR1 accumulated in the cellular fraction, and was very low or not detected in the medium, therefore the MFHR1 concentrations described were measured from cellular extracts in the whole study. Our results reveal a positive correlation across the three molecular levels DNA-copy number, mRNA-level and protein yield (Figure 1).

**Figure 1.**
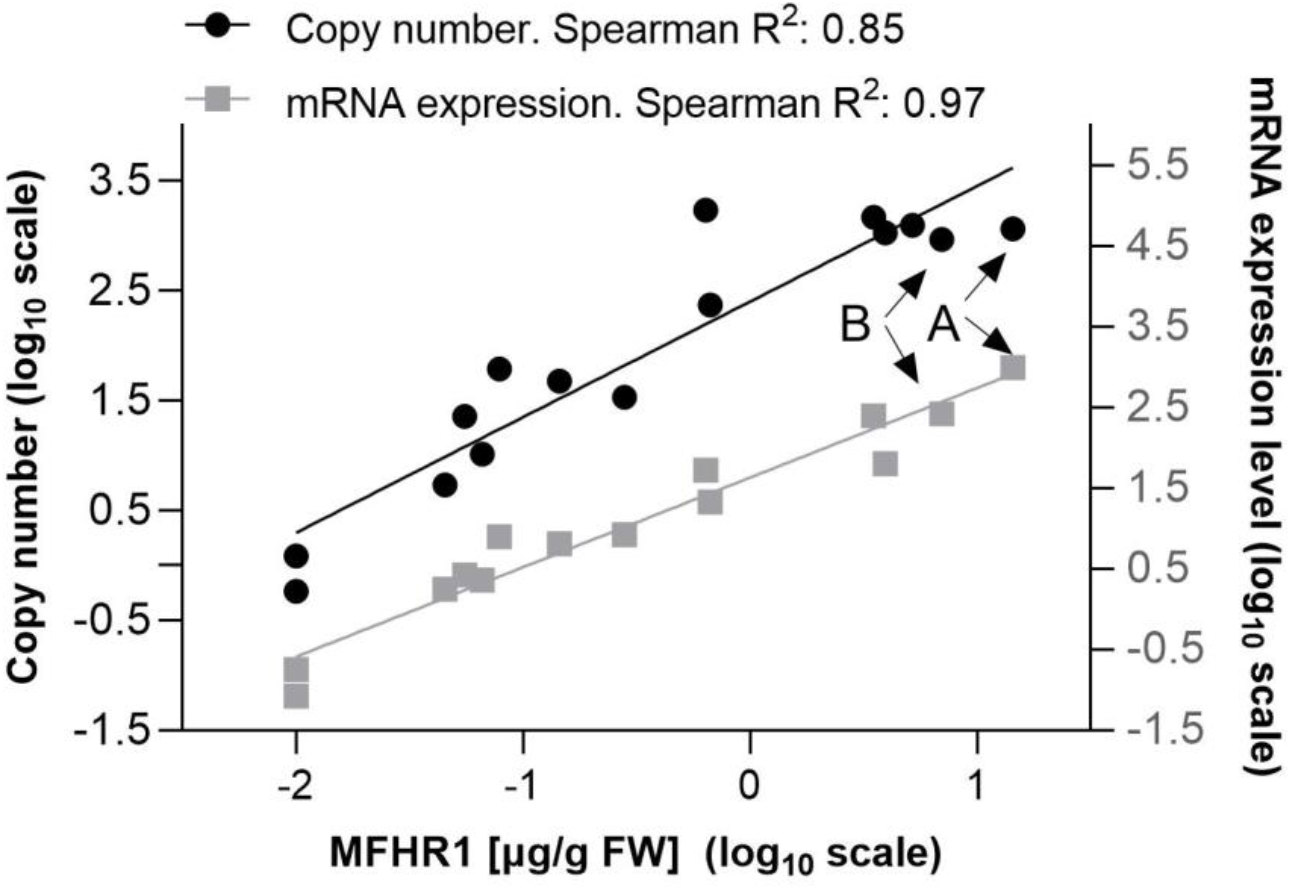
Positive correlation between three molecular levels: number of integrations of expression cassettes, mRNA expression level and recombinant protein production. The correlation was estimated with 14-15 MFHR1 producer lines, which include low and high producers. The experiment was performed one year after moss transfection. Linear regression in logarithmic scale is shown. Lines A and B selected for the next experiments are marked with arrows. mRNA expression levels were determined by qRT-PCR. Samples were normalized using two reference genes (coding for EF1-α and 60S ribosomal protein L21) and the line with the lowest expression level was used as calibrator. Number of integrations of expression cassettes was determined by qPCR. Samples were normalized with the single copy gene *PpCLF* (Pp3c22_22940V3.1).

According to previous studies protein and mRNA levels do not follow a normal distribution; therefore, the Spearman correlation was considered more suitable than the Pearson correlation (Maier et al., 2009). Accordingly, in our study mRNA expression and copy number of integrated expression cassettes could explain around 90% of the MFHR1 level variation (Figure 1). Many regulatory mechanisms could explain the remaining 10% such as DNA positional effects, methylation, miRNA and protein stability (Khraiwesh et al., 2010; Vogel and Marcotte, 2012; Myhre et al., 2013; Payne, 2015).

Although the correlation between protein and mRNA is weak or insignificant when groups of proteins or the proteome are analyzed (R^2^ 0.3-0.89) (Maier et al., 2009; Schwanhäusser et al., 2011), a better correlation was reported for individual proteins which are essential for growth, such as those implied in energy metabolism (Nie et al., 2006). This is congruent with our results, since MFHR1 expression is driven by a native *actin* gene promoter *(PpAct5)* (Top et al., 2019). Actins, essential components of the cytoskeleton, are ubiquitous proteins in eukaryotic cells and play important roles in cell division and extension, among others (Szymanski and Staiger, 2018; Wu and Bezanilla, 2018).

For further analyses we selected the lines with the highest productivity for each MFHR1 variant, referred from now on as line A and line B. At shaken-flask scale, line A (MFHR1^V62^) produces approx. 20-30 µg MFHR1/g fresh weight (FW) and is further referred to as the high-producer line, while line B (MFHR1^I62^) shows a lower productivity of up to 10 µg MFHR1/g FW. Recombinant protein production was scaled up to 5 L stirred tank bioreactors, and batch, fed-batch and semi-continuous operation modes were tested in order to increase protein productivity and yield. For the sake of clarity, in this study we referred to specific productivity as the amount of recombinant protein per gram fresh weight biomass, recombinant protein specific production rate as g recombinant protein /(g biomass d), while yield is the mass of accumulated recombinant protein during the whole process.

### 3.2 Mathematical modelling and effect of the bioreactor operating mode on recombinant protein productivity

Although model-based design aiming at process control and optimization can be very attractive for the biopharmaceutical industry to reduce costs and make data-driven decisions, it has been less implemented in plant-based expression systems (Craven et al., 2013). Bioreactor operation mode is one of the variables that needs to be optimized to increase product yield and to guarantee the reproducibility of the results. The choice of the operation mode in plants is host species-specific and depends on the expression system (constitutive or inducible promoter, intracellular or secreted protein) (Huang and McDonald, 2012). We evaluated different operation modes aiming at increasing the specific MFHR1 productivity to facilitate the purification of the protein from the cellular fraction. Furthermore, we used an unstructured kinetic model based on equations 1-11 described in the materials and methods section, which considers just the macroscopic variables, and tested its accuracy to predict recombinant protein production in moss suspension cultures in photobioreactors operated in batch, fed-batch and semi-continuous modes, respectively. The growth kinetic model, which describes the dynamics of biomass growth, MHR1 accumulation and nitrate consumption, was implemented with line B. The kinetic parameters were estimated using differential evolution algorithm implemented in SciPy (Python) (Table 1).

**Table 1.** Estimated kinetic parameters for batch, fed-batch and semi-continuous model of an MFHR1 producer line (Line B, MFHR1^I62^). r_X,max_: maximum specific growth rate. k_N_ and k_i_: saturation constants of the limiting substrates, nitrate and light, respectively. y_X,N_: biomass yield coefficient. r_p,max_: maximum specific recombinant protein production rate. k_p_: nitrate saturation constant for recombinant protein synthesis. k_pd_: specific protein degradation rate. s_X_: cell absorption cross section. r_R_: bioreactor cylinder radius. l_0_: incident light. NA: not-applicable.

|  | Batch<br>(parental line) | Batch | Semi-continuous |
| --- | --- | --- | --- |
| $r_{x,max} [\text{d}^{-1}]$ | 0.699 | 0.699 | 0.647 |
| $k_N [\text{mg/L}]$ | 3.58 | 4.41 | 3.91 |
| $k_i [\mu\text{mol}/(\text{m}^2 \text{ s})]$ | 15.01 | 15.29 | 16.86 |
| $y_{X,N} [\text{g/g}]$ | 9.615 | 7.042 | 5.050 |
| $r_{p,max} [\text{mg}/(\text{g d})]$ | NA | 0.327 | 0.236 |
| $k_p [\text{mg/L}]$ | NA | 1.010 | 1.545 |
| $k_{pd} [\text{L}/(\text{g d})]$ | NA | 0.0801 | 0.0801 |
| $\sigma_X [\text{m}^2/\text{g}]$ | 0.021 | 0.049 | 0.044 |
| $r [\text{cm}]$ | 8.75 | 8.75 | 8.75 |
| $I_0 [\mu\text{mol}/(\text{m}^2 \text{ s})]$ | 350 | 350 | 350 |

All bioreactors shared the same basic operation conditions: 0.3 vvm air with 2% CO_2_, 500 rpm stirring, and light intensity: 160 µmol/m^2^s (day 0 – 2), from day 2: 350 µmol/m^2^s. The batch run took eight days to reach stationary phase and MFHR1^-^specific productivity reached ∼ 65 ± 11.8 µg/g FW (Table 2) at the end of the exponential growth phase and decreased steeply afterwards. Furthermore, growth cessation coincided with the complete consumption of nitrate, therefore we considered it in addition to light as a limiting substrate in the dynamic model.

**Table 2.** Recombinant protein (MFHR1) productivity in several operation modes in moss bioreactors.

|  | **Batch** | **Fed-batch** | **Semi-continuous** |
| --- | --- | --- | --- |
| Time to achieve highest MFHR1 productivity [d] | 8 | 8 | 6 - 11 |
| Highest specific MFHR1 productivity [$\mu\text{g/g FW}$] | $65 \pm 11.8$ | $77.6 \pm 9.1$ | $57.6 \pm 7.1$ |
| Total MFHR1 [mg] | $10 \pm 1.7$ | $13.4 \pm 2.9$ | $6.6 \pm 0.6$ |
| Highest biomass density [g/L DW] | $3.39 \pm 0.06$ | $4.4 \pm 0.2$ | $1.67 \pm 0.04$ |
| Biomass density [g/L DW] at day of highest protein productivity | $3.16 \pm 0.03$ | $3.5 \pm 0.44$ | $1.2 \pm 0.05$ |

Initially, the parental line *(Δxt/ft)* was cultivated in batch operating mode and the proposed model was fitted to experimental data to predict biomass growth and nitrate uptake rates. Kinetic parameters of the parental line (Table 1, Supplementary Figure S1) allowed us to narrow the bounds to estimate the parameters of the moss transgenic line B. The maximum specific growth rate (r_x,max_) of the parental line was 0.699 d^-1^ (doubling time approx. 1 day) and the biomass yield was 9.6 g biomass/g nitrate (Table1). Our model fits accurately with the experimental data in batch operation until the beginning of the stationary phase, and it is able to predict concentration of product, nitrate and biomass (Figure 2A). The maximum specific growth rate of MFHR1-producing line B was almost identical to the parental line, but biomass yield was lower with 7.14 g biomass/g nitrate and maximal specific MFHR1 production rate (r_pmax_) was 0.32 mg protein/g biomass d (Table 1).

**Figure 2.**
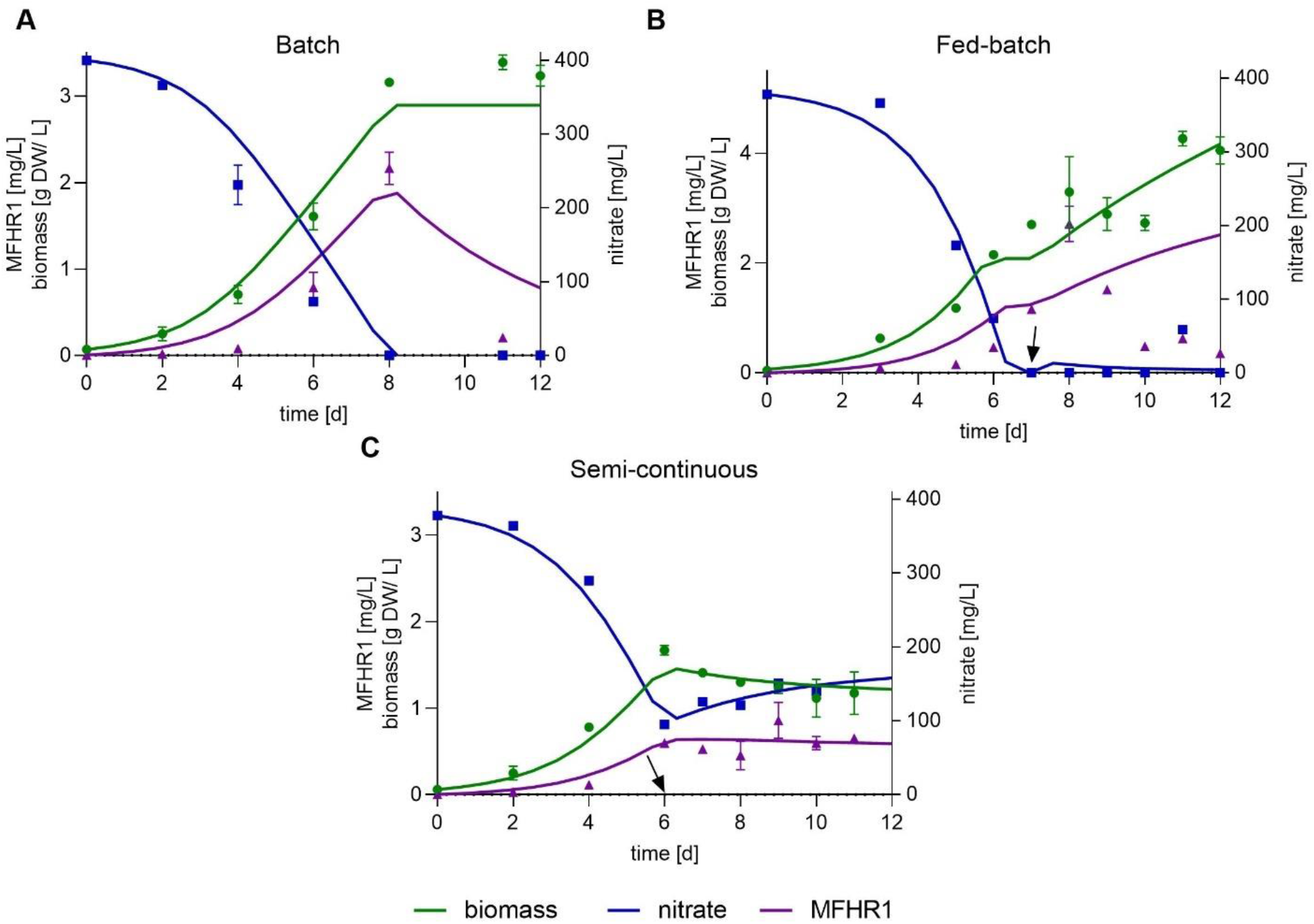
Mathematical modelling and simulation to describe the growth kinetics of the transgenic moss line B and the production of the recombinant protein MFHR1. Parameters were estimated using differential evolution (DE) implemented in Scipy in Python 3.6. The solid lines represent modelled data; experimental data are shown with symbols (**A**) Batch mode, (**B**) Fed-batch mode, (**C**) Semi-continuous mode. This operation was conducted between day 6 and 11. Data represent mean ± standard deviations (SD) from two measurements. The starting day of fed-batch and semi-continuous operations are marked by arrows.

To achieve a higher biomass and subsequent higher MFHR1 yield, we fed our culture with nitrate (fed-batch). The kinetic parameters obtained in the batch mode were initially used to calculate the feeding flow rate (F) maximizing biomass productivity and F was set at 200 mL/d with a nitrate concentration of 2,500 mg NO_3_^-^/L. The bioreactor was initiated with 4 L, fed-batch operation started at day 7 and the process was ended at day 12. The experimental data obtained were used to estimate new parameters to fit the fed-batch mode accurately. The proposed model was able to predict the dynamic behavior of biomass and nitrate but not recombinant protein. We observed again a drop in MFHR1 production between day 9 and 10 before reaching stationary phase of growth although nitrate was not limiting in this case (Figure 2B).

The discrepancy between model prediction and experimental data might be due to the age of the tissue and a change in the metabolic activities of the cells. After approx. nine days, the extracellular medium started turning slightly brownish, which might be attributed to accumulation of polyphenolic compounds and oxidative stress (Halliwell, 2003; Wilken and Nikolov, 2012). Although maximum biomass concentration increased from 3 g dry weight (DW)/L to 4 g DW/L compared to batch bioreactor, specific protein productivity and total product in the cellular fraction was similar (around 77.6 ± 9.1 µg MFHR1/ g FW, Table 2). Kinetic parameters are not listed in Table 1, due to the discrepancy between model prediction and experimental data.

The semi-continuous bioreactor was run for 11 days under the same conditions used for the batch mode. From day 6 to 11, 2 L suspension culture were replaced daily with the same volume of fresh medium, which corresponds to a dilution rate (D) of 0.4 d^-1^. However, as this D was slightly higher than the apparent specific growth rate (0.39 d^-1^, calculated between day 6-11), washing out of the cells was likely to occur. Kinetic parameters are listed in Table 1. The proposed model was able to accurately predict biomass growth, nitrate uptake and protein production (Figure 2C). The specific protein productivity was the lowest compared with fed-batch and batch (around 50 µg MFHR1/g FW). In total, around 10 mg and 13 mg recombinant protein accumulated in the cellular fraction after 8 days in batch and fed-batch, respectively, while for the bioreactor in semi-continuous mode around 6 mg were produced within 11 days, including biomass removed during this operation (Table 2). This operation could not be prolonged beyond 2 weeks, because the moss filaments formed big aggregates (pellets). This resulted in a reduced specific growth rate, making it difficult to reach a stable state (Supplementary Figure S2). Growth in pellets was also observed at the end of batch or fed-batch operation mode. Dense pellet formation could cause limitations in light, CO_2_ and nutrient transfer, which can affect product formation. However, limitations in CO_2_ could not be demonstrated in moss pellets (Cerff and Posten, 2012b). Formation of large entanglements and pellets can be avoided or delayed with low initial light intensities, however it reduces the maximum specific growth rate (Cerff and Posten, 2012b). The effect on recombinant protein production has to be further studied.

The kinetic model was validated to assess its predictability under different conditions in the semi-continuous operation mode, using the same estimated kinetic parameters (Table 1). For this the dilution rate and the starting time of the semi-continuous operation were modified to allow higher biomass density to accumulate and prevent washing out of the cells. From day 7 to day 10, 1 L of the suspension culture was replaced daily with the same volume of fresh medium (D = 0.2 d^-1^). Under these operation conditions, around 8.2 mg MFHR1 accumulated in the cellular fraction, 36% more than with a higher dilution rate (D=0.4 d^-1^). The model fits well with experimental values for biomass, nitrate and recombinant protein concentration with a coefficient of determination (R^2^) around 0.96, 0.99 and 0.88, respectively (Supplementary Figure S1B). According to the results, the model shows an acceptable predictability using several operating conditions with different variables.

Altogether, the specific MFHR1 productivity and total MFHR1 produced was affected by the change in medium conditions induced by the operating mode. The total amount of recombinant protein was doubled by using fed-batch or batch compared to semi-continuous operation (D=0.4 d^-1^), although the maximum specific productivity increased by just 35%. Here, we propose an unstructured and non-segregated kinetic model of the moss bioreactor to produce recombinant proteins, in this case MFHR1, which can be used as a starting point to optimize biomass and recombinant protein production under different operation modes or for the bioprocess design. The model fits accurately with the experimental data in batch and semi-continuous operation, and is able to predict recombinant protein production, nitrate uptake and biomass growth.

### 3.3 Effect of auxin on the production of MFHR1 at shaken-flask scale

Auxin plays an important role in many plant processes as in protonema development, promoting the transition from chloronema to caulonema cells (Decker et al., 2006). Exogenous auxin addition to the moss bioreactor triggered a temporal increase in MFHR1 production of 25% within 24 h (Top et al., 2019). Therefore, the effect of auxin on recombinant protein production was further studied in order to establish a strategy to increase protein yield.

First we studied the effect of the synthetic auxin naphthalene acetic acid (NAA, 10 µM) on MFHR1-specific productivity in the high-producer line A compared to the effect of the auxin biosynthesis inhibitor L-kynurenine (L-Kyn, 10 µM and 100 µM) (He et al., 2011), and their respective solvent controls (0.00005% KOH and 0.01% DMSO), in shaken-flask scale for 7 and 18 days.

Addition of auxin (10 µM NAA) increased MFHR1 concentration by 470% at day 7 compared to the controls (Figure 3A). In contrast, the specific protein production remained constant with 10 µM L-Kyn and decreased with 100 µM L-Kyn by 110 and 580% at day 7 and 18, respectively, while in the controls MFHR1 concentration doubled between day 7 and 18 (Figure 3A).

**Figure 3.**
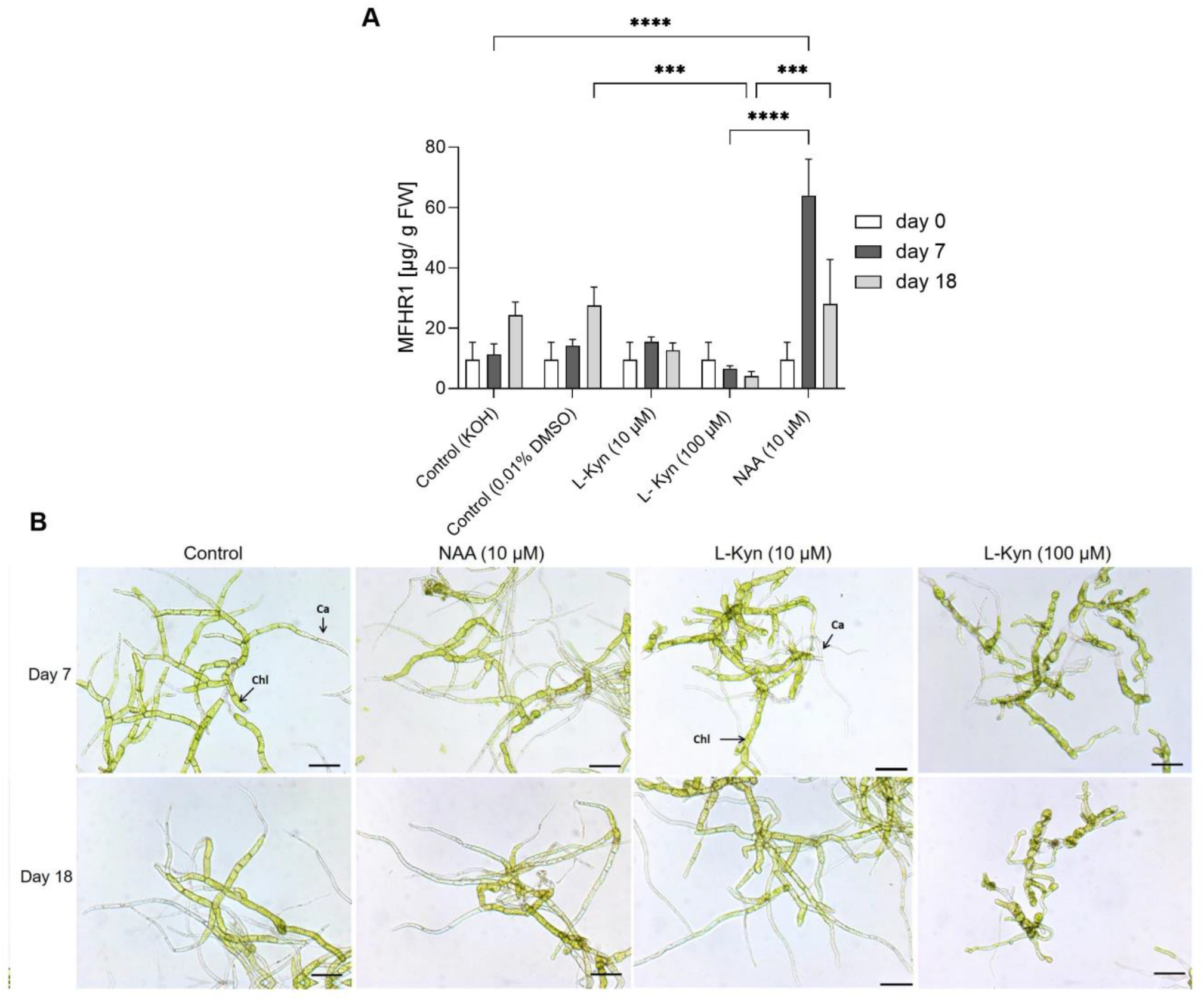
Exogenous auxin (NAA) enhanced recombinant protein production in Physcomitrella at shaken-flask scale, using transgenic moss line A. **(A)** Effect of exogenous auxin (NAA) and auxin biosynthesis inhibitor (L-Kyn) on MFHR1 production in Physcomitrella. NAA (10 µM) significantly enhanced recombinant protein specific productivity (P<0.0001 Two-way ANOVA, Tukey Post-hoc) compared with the control without NAA, while L-Kyn inhibited protein specific productivity significantly compared with the control (P= 0.0009, Two-way ANOVA, Tukey Post-hoc) and NAA treatment (P= 0.0007, Two-way ANOVA, Tukey Post-hoc) at day 18. Data represent mean ± standard deviations (SD) from three biological replicates **(B)** Representative light microscopic images of protonema tissue under different conditions growing in suspension in shaken flasks with NAA, L-Kyn and a control in standard medium Knop-ME. Scale bar: 100 µm.

The effects of auxin were also reflected by the morphological appearance of the tissues (Figure 3B). Compared to the control, NAA-treated suspensions showed accelerated caulonema development at day 7. In contrast, the addition of L-Kyn (10 and 100 µM) slowed down caulonema formation. Furthermore, the higher concentration of L-Kyn led to the formation of round, thick-walled brachycytes (Figure 3B), which are formed under stress or the addition of ABA (Decker et al., 2006). This observation supports a reduction of endogenous auxin levels by L-Kyn.

Next we studied the effect of different NAA concentrations (2 - 50 µM) at shaken-flask scale for 14 days. We did not find significant differences of specific recombinant protein productivity, although 50 µM NAA seemed to negatively affect protein production compared to 2-10 µM NAA (Supplementary Figure S3A). Likewise, we could not find significant differences in the growth index ((maximum biomass – initial biomass) /maximum biomass) upon different auxin concentration treatments at shaken-flask scale (Supplementary Figure S3B).

Auxin responses normally follow a bell-shaped dose-response curve. At small concentrations and up to an optimum, auxin induces cell division and elongation; beyond this maximum, opposite effects were observed in *Arabidopsis* shoots and cell suspension cultures of tobacco BY2, when inhibition of growth and bundling of actin occurs (Huang et al., 2017). According to previous studies, usually for high NAA concentrations (> 10 µM) the response is less pronounced (Waller et al., 2002; Huang et al., 2017). Therefore, we selected the concentration 10 µM for further studies in the bioreactor.

Our results confirm that exogenous auxin significantly enhances recombinant protein production in Physcomitrella (P<0.0001) and the inhibition of the endogenous synthesis of auxin by L-Kyn decreases the productivity of the protein of interest. Two different mechanisms can play a role in the MFHR1 production response to NAA treatment: tissue differentiation to the faster growing caulonema and an auxin signaling-dependent actin response, as the *actin5* promoter is driving the expression of our recombinant product (Top et al., 2019). The transition from chloronema to caulonema can be mediated by auxin or the quality of energy supply (Thelander et al., 2005; Decker et al., 2006). Caulonemal cells exhibit a faster elongation rate than chloronema (∼ 20 μm/h vs. ∼ 6 μm/h) (Menand et al., 2007). Optimal rates of tip growth are linked to this transition, which is characterized by non-stabilized actin cytoskeleton where actin genes are highly expressed. Furthermore, analysis of the transcriptome of caulonema and chloronema showed that all the processes related to tip growth are more active in caulonemal cells, such as cell-wall modification and regulation of cell size. An elongation of caulonemal cells can also result in enriched expression of certain actin genes (Ortiz-Ramírez et al., 2016). In addition, endoreduplication of the nuclear DNA was observed when caulonemal cells become older, and exogenous auxin led to a higher proportion of polyploid cells (Schween et al., 2003). Therefore, endopolyploidization can also contribute to the high specific recombinant protein productivity achieved upon auxin treatment.

### 3.4 Effect of sugar on the production of MFHR1 at shaken-flask scale

The quality of energy supply can influence tissue differentiation in Physcomitrella, e.g. exogenous glucose favors caulonema development. However, these cells are different from those induced by auxin, since they are shorter and “heavily pigmented” (Thelander et al., 2005). Therefore, we evaluated the influence of mixotrophic and heterotrophic conditions on the productivity of MFHR1.

Transgenic moss line A was cultivated under autotrophic, mixotrophic (1% sucrose and 50-70 µmol/m^2^s light, photoperiod 16/8), and heterotrophic conditions (1% sucrose and darkness). Protonema suspension cultures were initially adapted to each condition during two weeks before the experiment. Interestingly, sucrose had a negative impact on recombinant protein production. The specific protein production at day 7 was similar between autotrophic and mixotrophic conditions, however, it decreased to zero at day 18 under the latter (Figure 4A). Furthermore, recombinant protein concentration was almost negligible under heterotrophic conditions.

**Figure 4.**
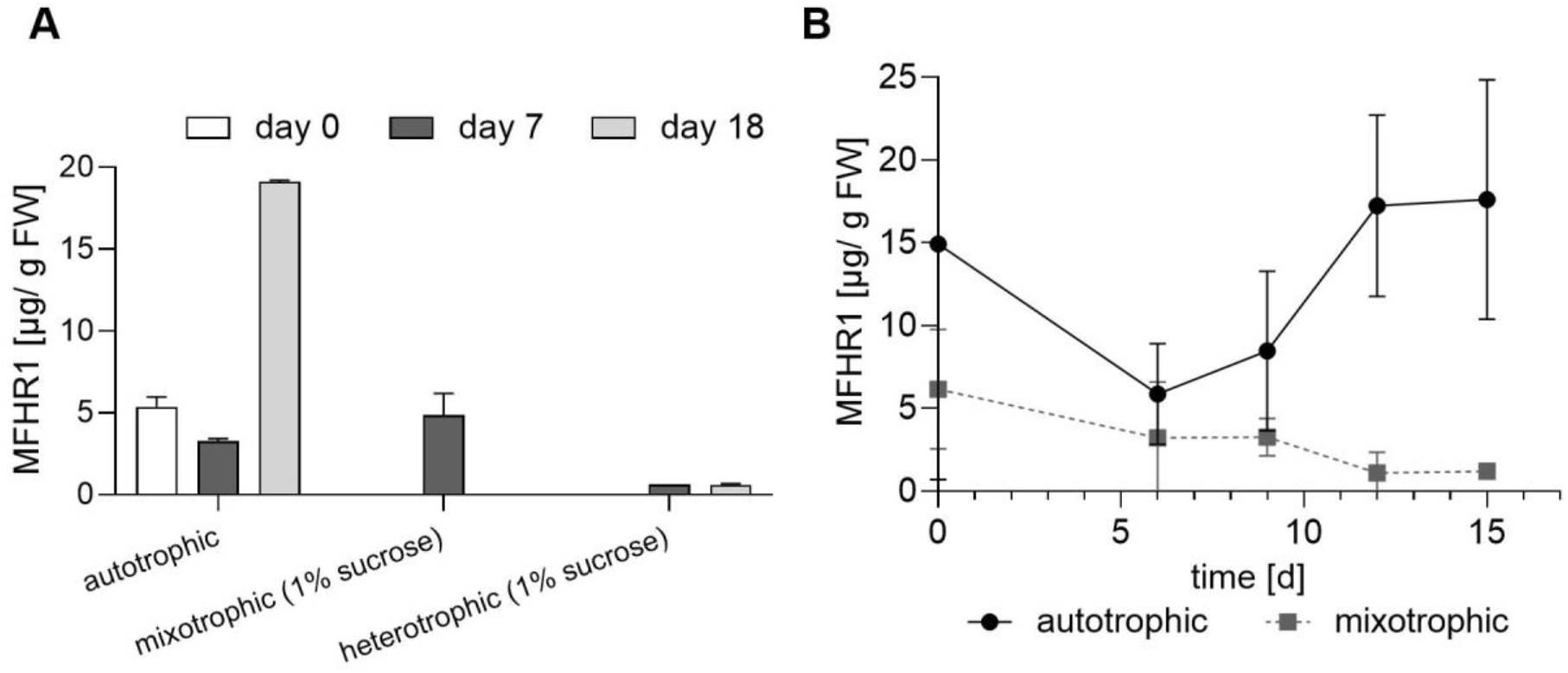
Mixotrophic and heterotrophic conditions negatively affect the production of MFHR1 **(A)** Specific MFHR1 productivity at shaken-flask scale under autotrophic (50-70 µmol/m^2^s light, photoperiod 16/8), mixotrophic (1% sucrose and 50-70 µmol/m^2^s light, photoperiod 16/8), and heterotrophic conditions (1% sucrose and darkness) at day 7 and 14 (transgenic line A). **(B)** Production kinetics of MFHR1 under autotrophic and mixotrophic conditions (1% glucose) in moss line A at shaken-flask scale. Data represent mean values ± SD from three biological replicates.

To analyze if the effect of sugar was dependent on the sugar type, we also tested 1% glucose for 15 days. Specific protein production decreased during the whole kinetics under mixotrophic as opposed to autotrophic conditions where protein specific productivity was ∼ 15-fold the productivity achieved with 1% glucose at day 15 (Figure 4B). There was no significant difference in terms of growth between medium supplemented with 1% glucose and the standard medium during 15 days at shaken-flask scale and nitrate was not limiting (Supplementary Figure S4).

### 3.5 Effect of auxin on the production of MFHR1 in 5 L bioreactor under different operating conditions

Subsequently, the effect of auxin supplementation was studied in 5 L stirred tank bioreactors operated in different conditions. Transgenic moss lines A and B were cultivated semi-continuously under the same conditions described above and 10 µM NAA were added daily from day 6 to keep the concentration stable (Figure 5). To explore a putative auxin influence on actin gene expression, a kinetic expression profile of the MFHR1 transgene upon auxin treatment was generated by quantitative real time PCR (qRT-PCR) from NAA-treated bioreactor samples at different time points. The expression of MFHR1 was compared to the auxin-responsive gene *PpIAA1A* and the actin gene *PpAct7*.

**Figure 5.**
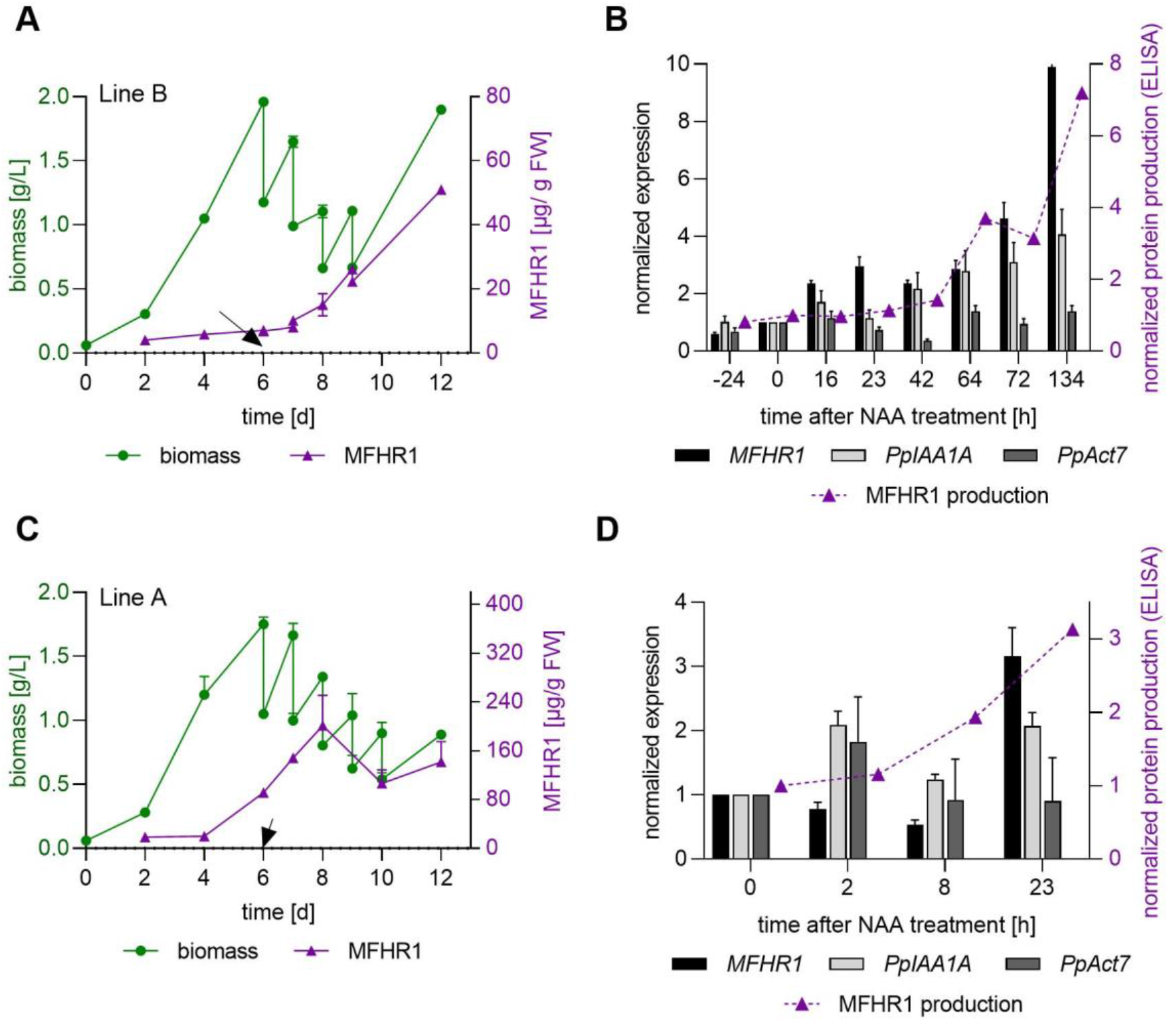
*MFHR1* transgene is upregulated during the whole growth kinetics, however, it is not rapidly induced upon auxin treatment. **(A), (C)**. Biomass and specific MFHR1 production in a semi-continuous bioreactor process with 0.4 d^-1^ dilution rate. Semi-continuous operation started at day 6 and 10 µM NAA were added. NAA was added daily to keep the concentration stable. **(B), (D)**. Expression of *Aux/IAA* gene (*PpIAA1A*: Pp3C8_14720V1.1), *PpAct7* (Pp3C3_33410V3.1) and *MFHR1* determined by RT-qPCR in 10 µM NAA-treated bioreactor samples. Data were normalized using two reference genes (coding for EF1-α and 60S ribosomal protein L21) and a time point immediately before NAA addition was used as calibrator. A comparison between the protein accumulation measured by ELISA and the expression of mRNA by RT-qPCR is included. Protein production quantified by ELISA was normalized with the amount produced immediately before NAA addition. Mean ± standard deviations (SD) from three technical replicates are shown. **(B), (D)** Data obtained from the bioreactor shown in **(A), (C)**. The arrow indicates the first addition of NAA.

Aux/IAA proteins have an essential role in auxin response and gene regulation. Physcomitrella has three genes (P*pIAA1A, PpIAA1B, PpIAA2*) encoding these proteins, and a triple knockout of the genes rendered the plants completely insensitive to auxin (Lavy et al., 2016). Therefore, *PpIAA1A* (Pp3C8_14720V3.1), which is induced upon auxin treatment (Prigge et al., 2010), was chosen as a positive control during NAA treatment in the bioreactor.

As the expression of *MFHR1* is driven by a *PpAct5* promoter, we aimed to test the response of *MFHR1* and *actin* gene expression, respectively, to auxin treatment. However, the Physcomitrella genome contains a duplication (Pp3c10_17070V3.1, *PpAct5b*) of *PpAct5* (Pp3C10_17080V3.1, *PpAct5a*) with a different promoter sequence but almost identical coding sequences. Therefore, *PpAct5* mRNA was not eligible for specific amplification. As the *PpAct5a* UTRs are present in our transgene construct, these sequences could not be used as a target for the analysis of *PpAct5a* expression in MFHR1 expressing lines either. A multigene family with different expression patterns and functions encodes actin proteins. *P. patens* has eight *actin* genes with highly conserved coding regions. Although all actin genes are expressed in protonema, *PpAct5a, PpAct7a* (Pp3c3_33410V3.1), and *PpAct7b* (Pp3c3_33440V3.1) are highly expressed (Supplementary Figure S5), according to the expression data obtained from PEATmoss (Fernandez-Pozo et al., 2020). Moreover, according to a distance-based dendrogram (Supplementary Figure S6), *PpAct5* and *PpAct7* are closely related and the expression levels of *PpAct5a* and *PpAct7a* are similar and higher than *PpAct7b* (previously called *PpAct3*) (Weise et al., 2006). Therefore, the expression level of *PpAct7a* was evaluated in the MFHR1-expressing moss line under auxin treatment.

*PpAct7a* expression was relatively stable during the bioreactor run suggesting that this gene is not regulated by auxin (Figure 5B), as opposed to *Arabidopsis thaliana Act7*, which is the only *Arabidopsis actin* gene strongly responding to auxin (McDowell et al., 1996). However, the *Aux/IAA* gene and *MFHR1* transgene were upregulated during the whole kinetics (Figure 5B). Interestingly, the *MFHR1* transgene was upregulated during the first 64 h in a similar way to the *Aux/IAA* gene, whilst a further increase in transgene expression compared to *Aux/IAA* was observed after 72 h when the semi-continuous operation was stopped for 2 days to let biomass increase. Thus, MFHR1 RNA level-increase coincided with increased biomass and MFHR1 protein accumulation (Figure 5A, B).

The transgene was upregulated within 16 h after NAA addition (Figure 5B). To further unravel the kinetic response of *MFHR1* to NAA, the expression level was analyzed within 2-23 h after NAA addition, using the MFHR1 high-producer line A (Figure 5C, D). The expression level of the transgene was constant up to 8 h after NAA addition, and clearly upregulated after 23 h while the *Aux/IAA* gene was induced already within 2h (Figure 5D), which suggests that the effect of auxin on *PpAct5* promoter-driven expression is indirect.

Although *PpAct5* was slowly induced upon auxin treatment, our results can be explained by a link between auxin signaling and actin. The debundling of actin is necessary for an efficient cellular response to auxin and promotion of auxin transport (Nick et al., 2009; Guillory and Bonhomme, 2021). Both, the natural and the artificial auxins, indole-3-acetic acid (IAA) and NAA, respectively, induced debundling of actin filaments. In *Arabidopsis*, overexpression of the actin-binding domain of plant fimbrin (GFP-FABD2) decreased auxin transport significantly due to reduced actin dynamics (Zaban et al., 2013). Furthermore, the overexpression of the same domain led to desensitization of auxin response in tobacco BY2 cell suspension cultures. These observations reveal that the role of actin in auxin signaling is not just a structural effect, since actin is involved in polar auxin transport (Bennett et al., 2014; Huang et al., 2017).

We observed a high increase in MFHR1 production in the bioreactor with NAA after the semi-continuous operation was ceased and the bioreactor was run in a batch phase for two days before ending the whole process (Figure 5A). This observation suggests that biomass concentration plays an important role in MFHR1 production, which concurs with our phenomenological model. Therefore, batch, fed-batch and semi-continuous bioreactors with lower dilution rates and auxin supplementation were evaluated. NAA (10 µM) was added at day 3 and 4 in the batch bioreactor process, reaching 823 µg MFHR1/g FW (150 mg MFHR1 accumulated over 7 days) and 200 µg MFHR1/g FW with line A and B, respectively (Figure 6). Compared to batch bioreactor without exogenous NAA, this treatment led to an approx. 230% increment in MFHR1 productivity with line B (Figure 6B). Product formation decreased at day 8, which could be attributed to nitrate limitation. With these conditions and auxin supplementation, we reached 8-fold the productivity achieved in an earlier study under semi-continuous operation mode for line A (Top et al., 2019) and 2.7-fold the specific productivity achieved under the same conditions but with auxin supplementation since day 6 (Figure 5C). During the bioreactor run with exogenous NAA (day 3), caulonemal cells started developing from day 2, even before auxin addition, and became more evident from day 4. Big pellets were evident from day 8 (Supplementary Figure S7). Auxins are mainly extracellular hormones (Reutter et al, 1998); endogenous auxin levels and cell density can influence the development of caulonema already at day 2. In *Funaria hygrometrica*, the effect of auxin is highly dependent on the cell density, and the manipulation of both can lead to cultures enriched predominantly with caulonema. At low cell densities (< 200 mg/L) basal caulonema development depends on endogenous auxin levels, and below 50 mg/L approx. 10% of cells starts differentiation (Johri, 2020).

**Figure 6.**
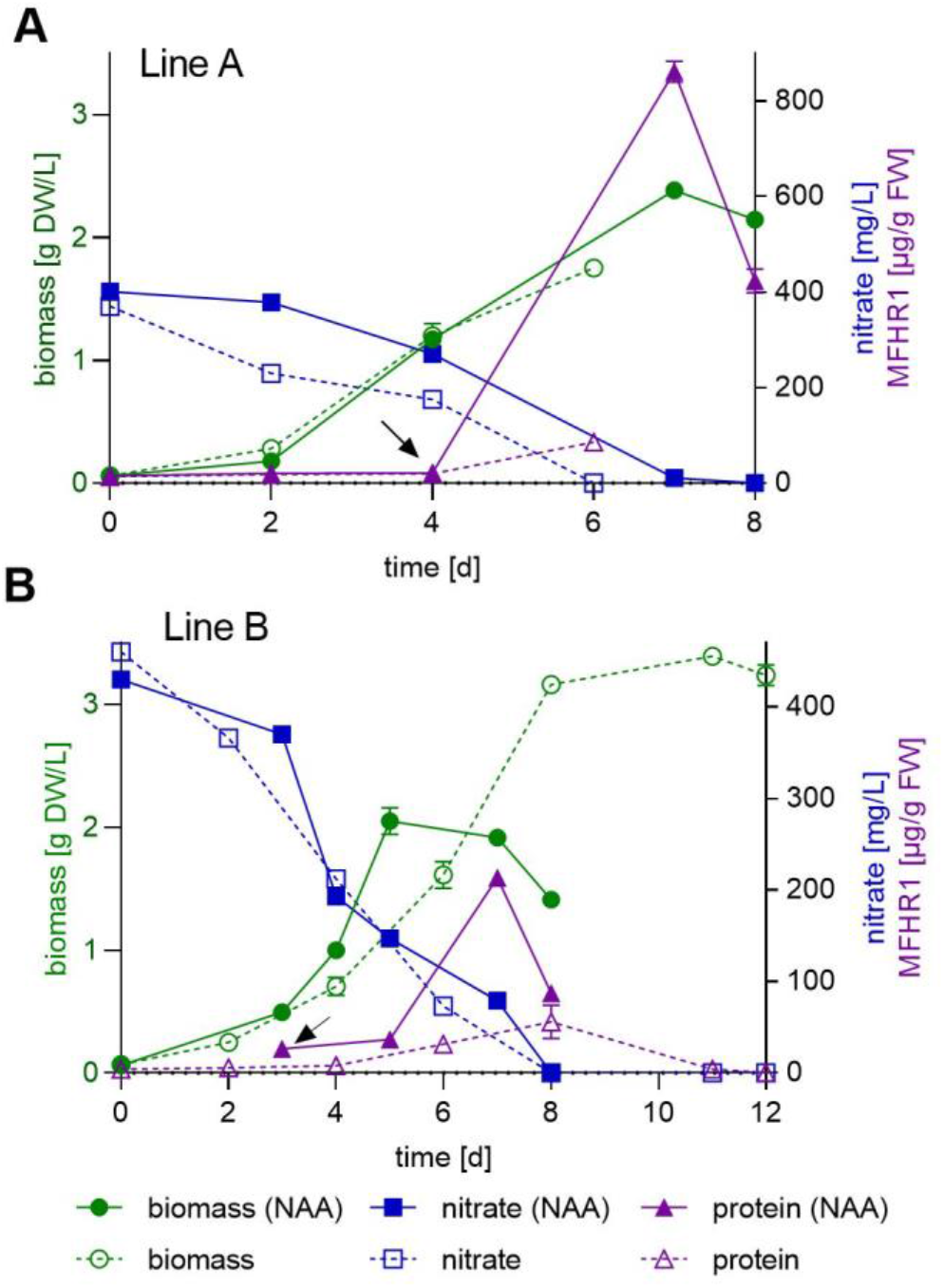
Exogenous auxin enhanced recombinant protein specific productivity in 5 L bioreactor operated in batch. Comparison of biomass, nitrate consumption and MFHR1 concentration between batch bioreactor run with (full lines) and without exogenous NAA addition (dashed lines). NAA (10 µM) was added at day 3 (**B**) or 4 (**A**). Data represent mean values ± SD from two independent measurements. The arrow marks the time point of NAA addition.

As the steep increase in MFHR1 accumulation was not immediate (Figure 5A), different time points for the beginning of NAA treatment in fed batch and semi-continuous processes were evaluated using line B (Figure 7). NAA (10 µM) was added at day 3, 5 or 7, respectively. Fed-batch operation started at day 7 by adding 200 mL/d (5x KNOP/ME), which corresponds to a concentration of 2,500 mg NO_3_^-^ /L. Semi-continuous operation started at day 7, and the dilution rate was reduced from 0.4 to 0.2 d^-1^ (1 liter culture was replaced daily instead of 2 liter) to allow the biomass to increase (Figure 7). Under these conditions in semi-continuous mode, the highest specific protein productivity was between 134 and 178 µg MFHR1/g FW. However, washing out of the cells was not prevented by decreasing the dilution rate in the case of NAA supplementation. As semi-continuous operation was not the best mode to achieve a high MFHR1 specific productivity we abstained from evaluation of NAA supplementation at day 5 (Figure 7B).

**Figure 7.**
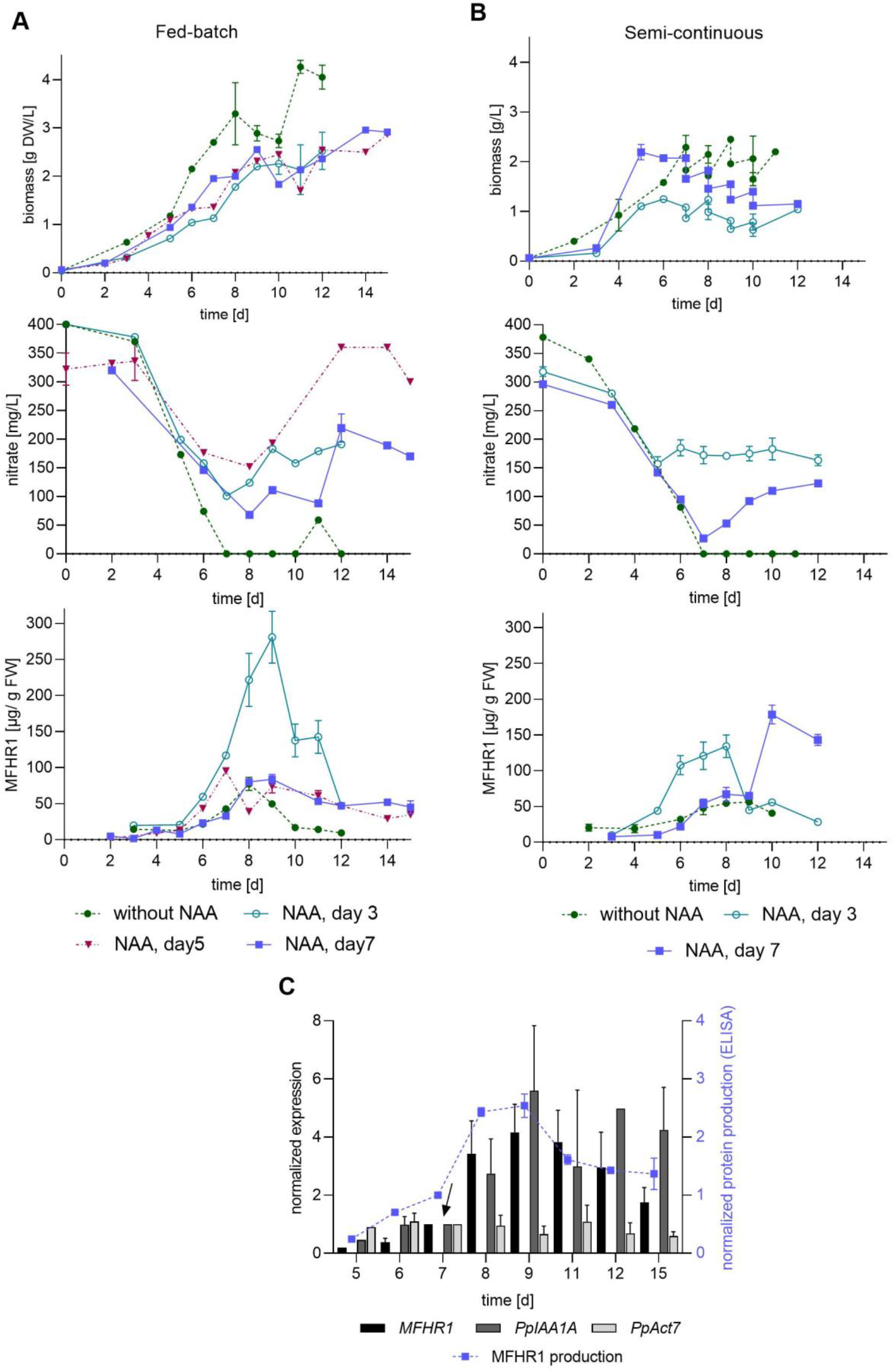
Effect of operation conditions and NAA addition time on recombinant protein production. **(A)** Fed-batch with and without NAA using transgenic moss line B. **(B)** Semi-continuous mode with and without NAA using line B. NAA was added at day 3, 5 and 7, respectively. **(C)** Expression of *Aux/IAA* gene (ppIAA1A: Pp3C8_14720V1.1), *PpAct7* (Pp3C3_33410V3.1) and *MFHR1* determined by qRT-PCR of 10 µM NAA-treated bioreactor samples. Data were normalized using two reference genes (coding for EF1-α and 60S ribosomal protein L21) and a point immediately before the addition of NAA (indicated by the arrow) was used as calibrator. A comparison between the protein production measured by ELISA and the expression of mRNA by RT-qPCR is included (data obtained from the bioreactor shown in (A), NAA added at day 7). Mean ± standard deviations (SD) from three technical replicates are shown.

In fed batch the time point of NAA addition was crucial. The highest specific recombinant protein productivity was obtained in fed-batch mode. When NAA was added at day 3 it reached up to 280 µg MFHR1/g FW, while only a third of this concentration was achieved when NAA was added later, at day 5 or 7 (Table 3, Figure 7A). The total amount of MFHR1 was doubled by the use of NAA from day 3 in fed-batch and the specific protein productivity increased by 260% and 230% compared to the fed-batch and batch operation without NAA, respectively (Tables 2, 3).

**Table 3.** Biomass and recombinant protein (MFHR1) production of line B under different operation conditions with exogenous auxin NAA (10 µM).

|  | **Batch** | **Fed-Batch** | **Fed-Batch** | **Fed-Batch** | **Semi-continuous** | **Semi-continuous** |
| --- | --- | --- | --- | --- | --- | --- |
| NAA (10 μM) treatment start [d] | 3 | 3 | 5 | 7 | 3 | 7 |
| Time to achieve highest productivity [d] | 7 | 9 | 7 | 9 | 8 | 10 |
| Total MFHR1 [mg] | 20.4<br>$\pm 0.6$ | 28.4<br>$\pm 3.6$ | 9.4<br>$\pm 1.2$ | 9.8<br>$\pm 0.8$ | 9.6<br>$\pm 1.1$ | 15.8<br>$\pm 1.2$ |
| Highest specific productivity [μg/g FW] | 213.2<br>$\pm 6.2$ | 280.9<br>$\pm 36.0$ | 95.5<br>$\pm 2.5$ | 83.8<br>$\pm 6.6$ | 134.3<br>$\pm 15.8$ | 178<br>$\pm 13.1$ |
| Highest biomass density [g/L DW] | 2.15<br>$\pm 0.11$ | 2.5<br>$\pm 0.22$ | 2.87<br>$\pm 0.08$ | 2.9<br>$\pm 0.01$ | 1.23<br>$\pm 0.19$ | 2.07<br>$\pm 0.03$ |
| Biomass density at day of highest productivity [g/L DW] | 1.91<br>$\pm 0.05$ | 2.2<br>$\pm 0.04$ | 1.4<br>$\pm 0.07$ | 2.55<br>$\pm 0.19$ | 1.23<br>$\pm 0.19$ | 1.39<br>$\pm 0.04$ |

Parameters of semi-continuous operation supplemented with auxin also influenced the product yield, which increased by 250% by decreasing the dilution rate, and delaying the starting day of the semi-continuous operation to allow higher biomass accumulation (Figure 5A, Figure 7B). These results suggest that cell density or tissue age play a role in protein production. The daily medium exchange can dilute metabolites involved in signaling, such as cytokinin and auxins that accumulated in the medium or influence the differentiation of the tissue, with a negative impact on recombinant protein production. However, this operation mode also avoids the steep decrease in MFHR1 production observed under batch or fed-batch conditions.

A drop in MFHR1 concentration was observed in fed-batch in each evaluated condition approx. after 8 days, similar to the behavior without auxin (Figure 7A), and is not related to nitrate limitation. A drop in protein accumulation might be due to a decrease in the expression of the transgene but also attributed to inefficient protein extraction, secretion of the protein to the extracellular medium or degradation. In order to further understand this decrease in protein concentration, we analyzed the transgene expression levels from fed-batch bioreactor samples treated with NAA at day 7. MFHR1 productivity quantified by ELISA was congruent with the transgene expression the first 3 days after NAA exposure (Figure 7C). The change in expression levels remained relatively stable from day 1 to 5 after NAA exposure, and decreased two days later, while MFHR1 productivity decreased after four days (Figure 7C). The expression patterns of the *MFHR1* transgene, *Aux/IAA* gene and *PpAct7* were similar to the results observed under semi-continuous operation (Figure 5B, 7C). Our results explain only partially the kinetics of MFHR1 concentration, as a lower rate of production caused by a lower expression of the transgene cannot be solely responsible for a decrease in concentration. The mechanisms involved in a drop of MFHR1 remain unclear. As the culture supernatant turns slightly brownish at the time where protein concentration decreases, we speculate that accumulation of polyphenolic compounds is a crucial factor. Moreover, this phenomenon occurred faster upon auxin treatment than in the control and could be a stress signal leading to accumulation and oxidation of these compounds and impairment of growth rate. During the extraction of the protein, covalent bonding between protein and phenolic compounds can occur and alter protein functionality and physico-chemical properties (Menkhaus et al., 2004), which might affect the estimation of MFHR1 concentration by ELISA. Likewise, auxin transport and response is inhibited when there is an accumulation of flavonoids (Mccurdy et al., 2001; Moody et al., 2021), which might explain the difficulty to increase protein production in the fed-batch mode with exogenous auxin even more.

In summary, we propose a phenomenological model to predict recombinant protein production in moss bioreactors operated in batch and semi-continuous operation, which can be used to optimize recombinant protein yield and biomass. This kinetic model can be applied to other plant cell suspension cultures to predict biopharmaceutical production, just by changing the terms of the energy source. Furthermore, exogenous auxin increased specific recombinant protein production by 470% in shaken flasks, and by up to 230% and 260% in moss bioreactors operated in batch and fed-batch mode, respectively. Under our conditions, the semi-continuous operation mode was inferior to batch or fed-batch for the production of MFHR1. A change in bioreactor operation mode along with NAA supplementation led to an increase of MFHR1 specific productivity up to 8-fold (0,82 mg MFHR1/g FW) compared to the values reported for line A (Top et al., 2019).

We conclude that *PpAct 7* and *PpAct 5* are not rapidly induced upon auxin treatment, which suggests an indirect response to auxin leading to upregulation of a transgene driven by the actin 5 promoter in Physcomitrella. The application day of auxin and the bioreactor operation mode are important factors to increase the recombinant protein specific productivity. In this case, MFHR1 production follows a mix-growth associated pattern. Therefore, a high growth rate and a high biomass density are essential.

We suggest that the auxin effect on biopharmaceutical production in moss bioreactors can be extrapolated to other plant-cell suspension cultures. Auxin stimulates the mitotic activity in tobacco-cell suspension cultures (Huang et al., 2017), which is advantageous to producing therapeutic proteins, when their expression is driven by a constitutive promoter. Although the medium of plant-cell suspension cultures usually includes cytokinins and auxins, the optimization of auxin concentrations is worth being considered, e.g., medium optimization of tobacco-cell suspension cultures producing the antibody M12 revealed a strong influence of the auxins IBA and 2,4-D (Vasilev et al., 2013).

The effect of auxin was not included in the phenomenological model due to lack of understanding of the relationship between auxin and the macroscopic variables biomass, nitrate and protein concentration. However, the data generated in this study will be the basis for implementing a data-driven model involving the effect of auxin in fed-batch bioreactors based on machine-learning approaches to further optimize recombinant protein production.

## 5. Author Contributions

NRM performed experiments and built the dynamic model. SS performed experiments at shaken-flask scale. NRM, JP, CP, RR, ELD conceived the study and wrote the manuscript. All authors read and approved the final manuscript.

## 7 Acknowledgements

We thank Andrés Felipe Posada-Moreno (Institute for Data Science in Mechanical Engineering, RWTH Aachen University, Aachen, Germany) for his kind assistance in parameter estimation of the kinetic model. We thank Prof. Peter Zipfel (Leibniz Institute for Natural Product Research and Infection Biology, Friedrich Schiller University, Jena, Germany) for providing us the polyclonal antibody anti-FH SCRs 1-4 and Anne Katrin Prowse for proof-reading of the article.

## Supplementary Material

**Supplementary Table S1.**
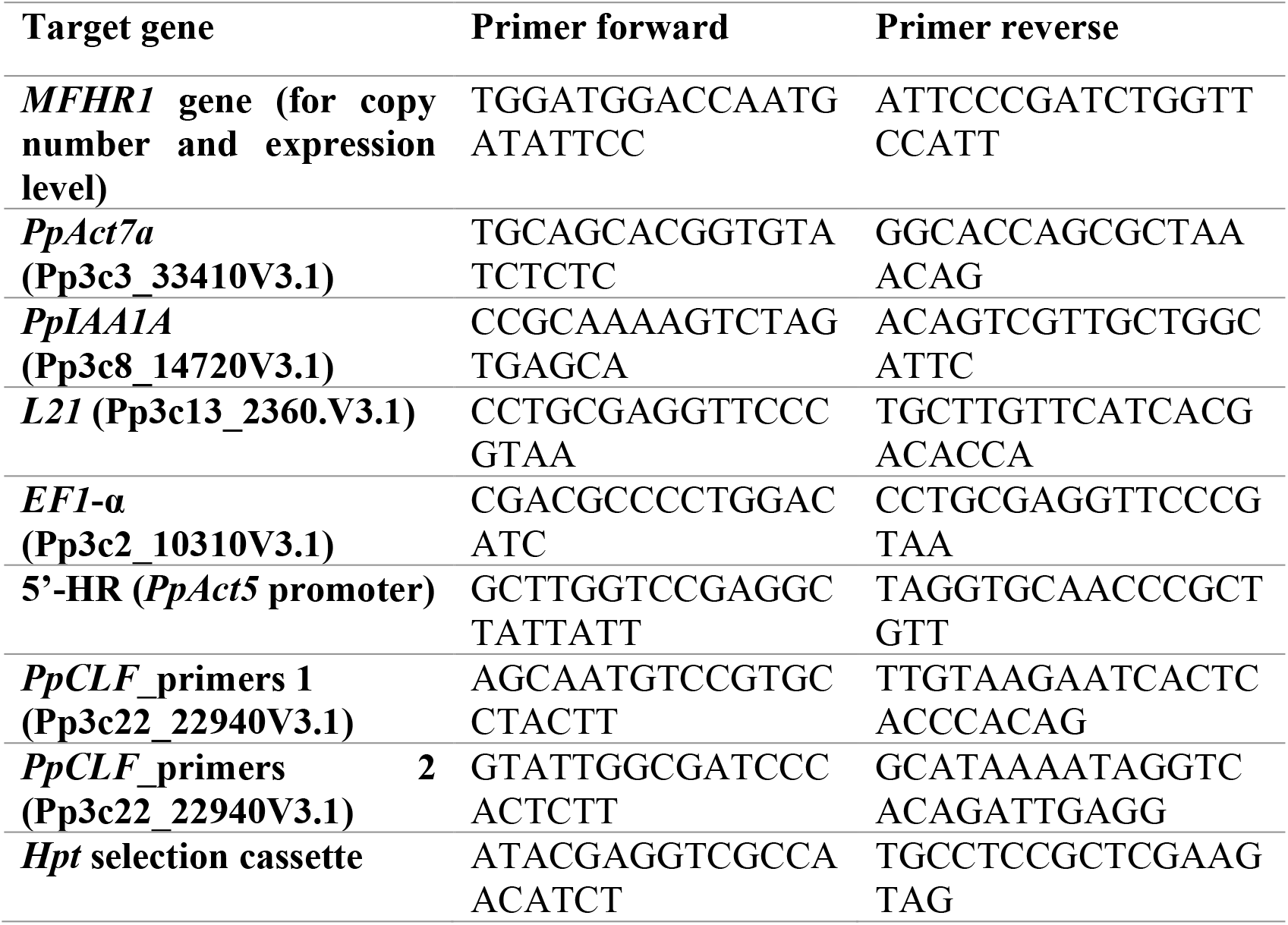
Primers used for qPCR and qRT-PCR to analyze number of expression cassettes integrated in the genome and expression levels of the transgene, the auxin responsive gene *PpIAA1A*, and *actin* gene *PpAct7a*. The gene model used for the prediction of the sequence is mentioned between brackets.

**Supplementary Figure S1.**
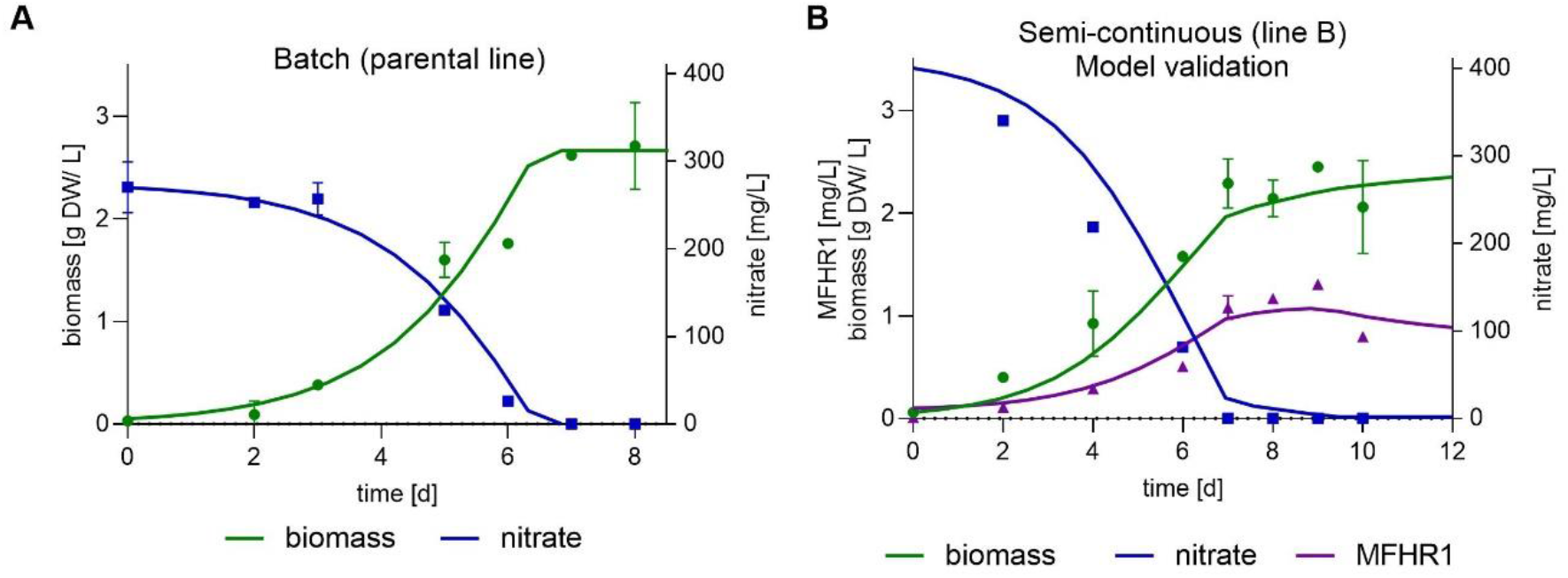
Mathematical modelling and simulation to describe the growth kinetics of Physcomitrella and the production of the recombinant protein MFHR1. (**A**) parental line (*Δxt/ft*) (**B**) Validation of the kinetic model using semi-continuous operation mode under different conditions. The operation started at day 7 with D= 0.2 d^-1^. Estimated parameters listed on table 1 were used. The solid lines represent modelled data; experimental data are shown with symbols. Parameters were estimated using differential evolution (DE) implemented in Scipy in Python 3.6. Data represent mean ± standard deviations (SD) from two measurements.

**Supplementary Figure S2.**
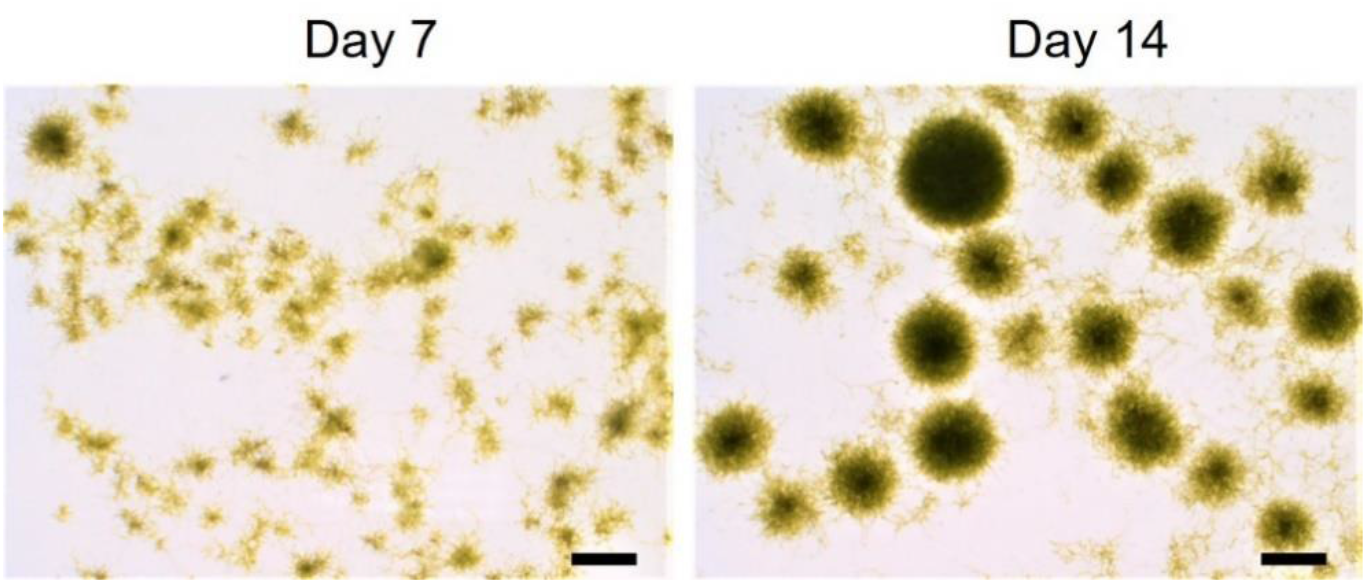
Light-microscopic photographs of moss-suspension cultures in stirred-tank bioreactor operated in semi-continuous mode. Big pellets are observed at the end of the operation. Scale bar: 1 mm

**Supplementary Figure S3.**
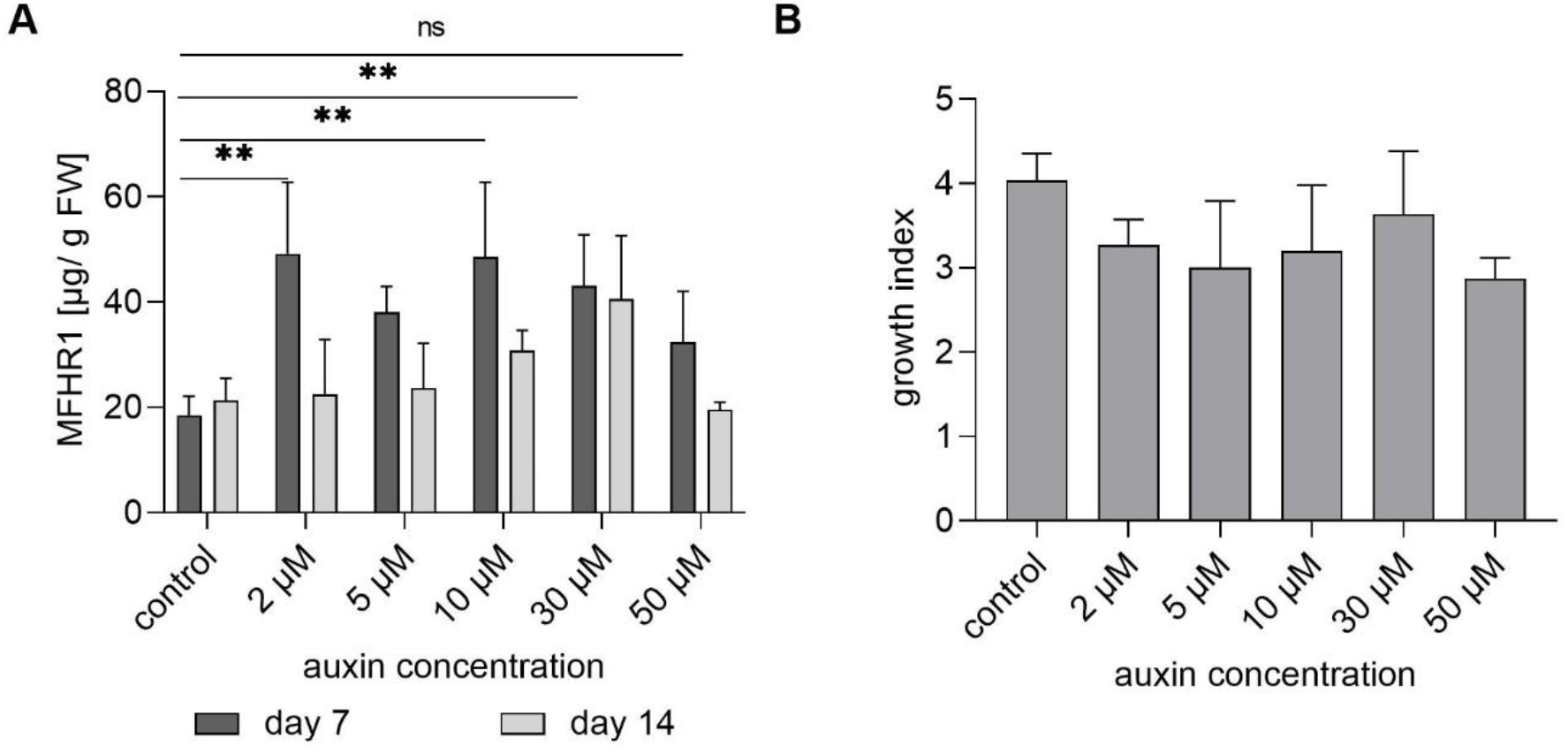
Effect of NAA concentration on specific MFHR1 production **(A)** There were no significant differences in MFHR1 productivity upon NAA treatment at different concentration (2-50 µM) at shaken-flask scale (One-way ANOVA, Bonferroni Post-hoc). Significant differences compared to the control without NAA are shown at day 7 (One-way ANOVA, Dunnet Post-hoc). Data represent mean values ± SD from three biological replicates. **(B)** Growth index of transgenic moss line A cultivated in shaken flasks under different NAA concentrations. Data represent mean values ± SD from three biological replicates.

**Supplementary Figure S4.**
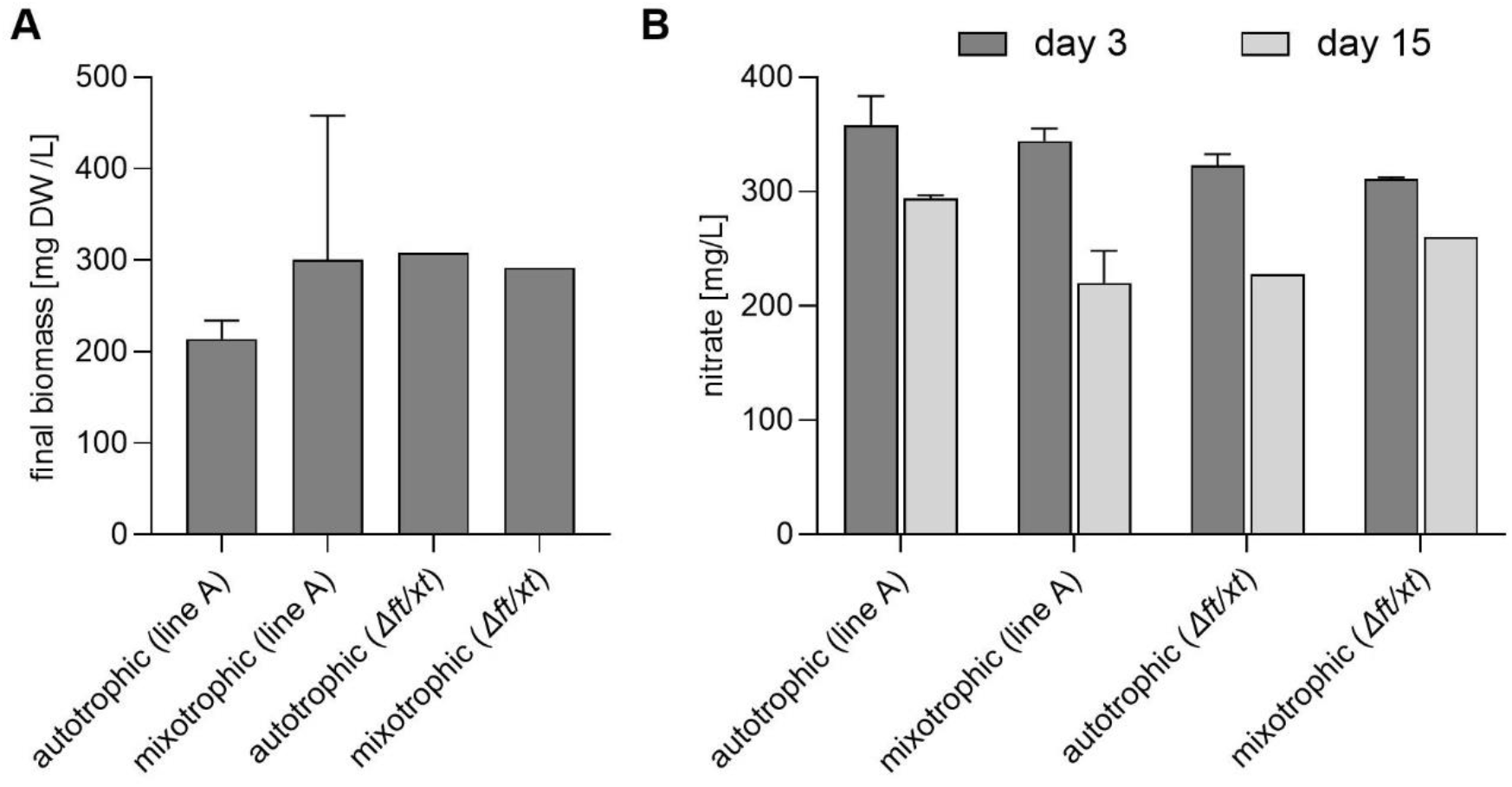
Effect of sugar addition on the growth of Physcomitrella. **(A)** Final biomass after 15 days. The initial cell density was 100 mg DW/L **(B)** Nitrate uptake at shaken flask scale under autotrophic and mixotrophic (1% sucrose and 50-70 µmol/m^2^s light, photoperiod 16/8 h) conditions. Moss transgenic line A and the parental line *Δ ft/xt* were used. Data represent mean values ± SD from three biological replicates.

**Supplementary Figure S5.**
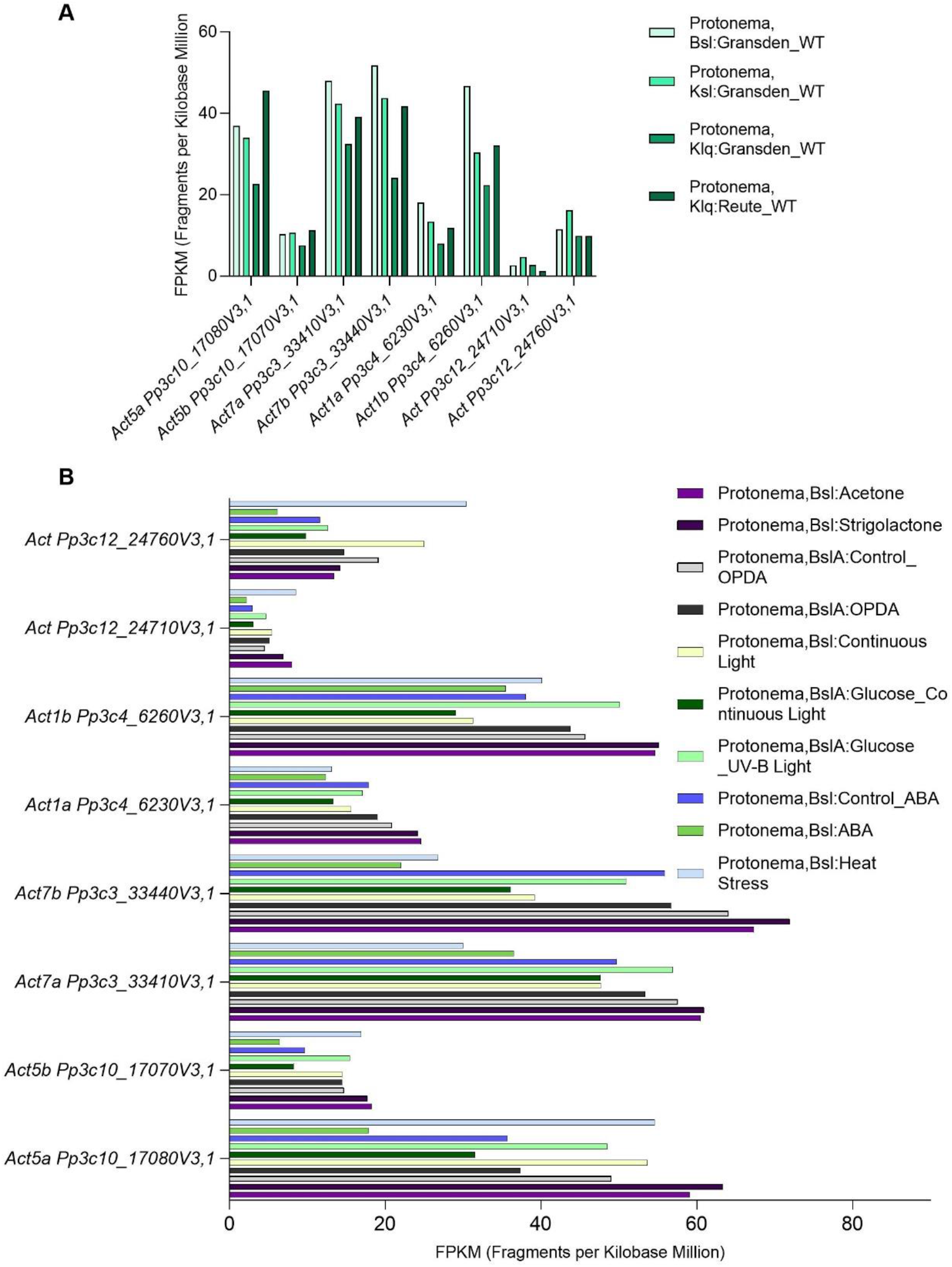
Actin genes expression levels in protonema of Physcomitrella obtained from PEATmoss (Fernandez-Pozo *et al*., 2020). (**A**) Expression in Reute and Gransden ecotypes. Dataset “RNAseq developmental stages” was used. (**B**) Expression in protonema under different conditions. Dataset “RNAseq protonema treatments” was used. The abbreviations describe the culture medium, Bsl = BCD solid, BslA = BCDA (ammonium), Klq = Knop liquid, Ksl = Knop solid.

**Supplementary Figure S6.**
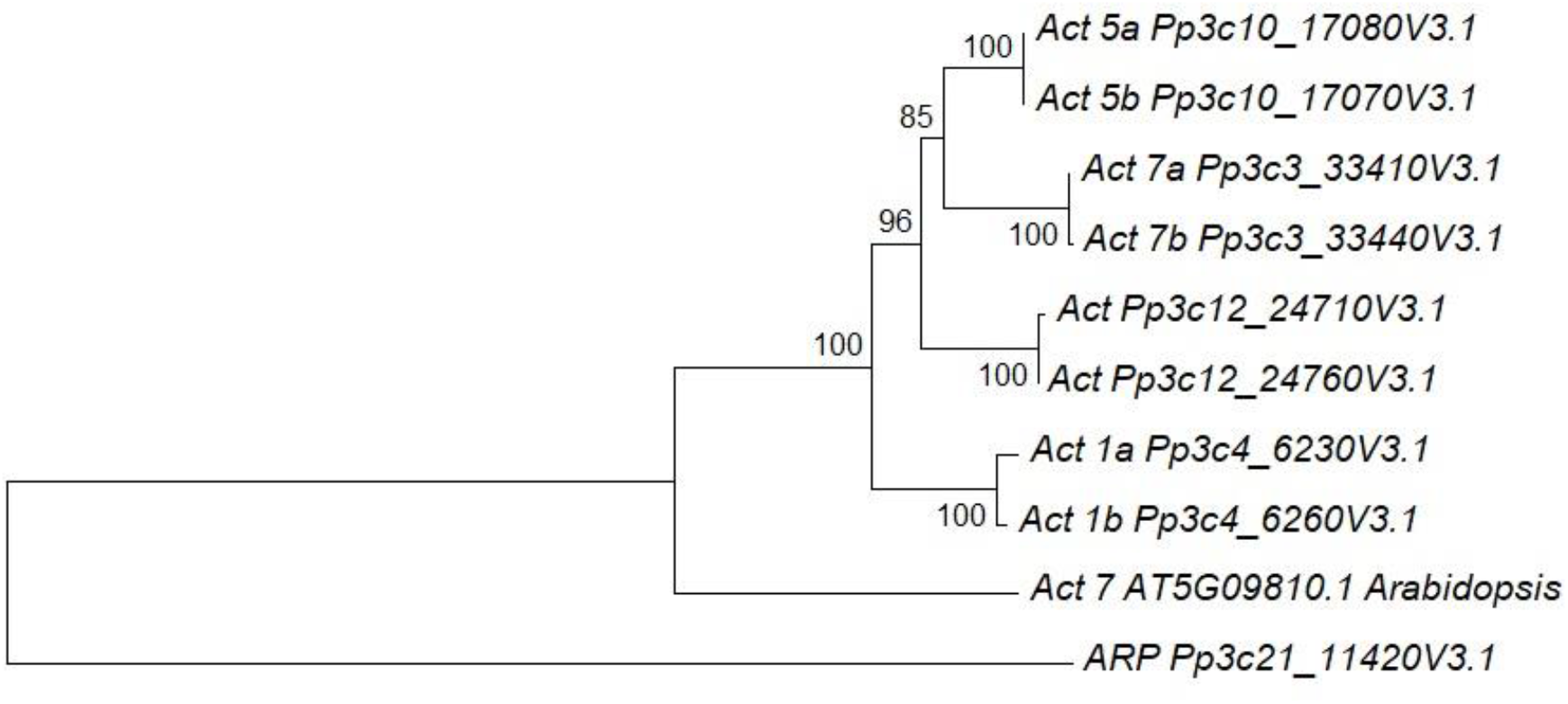
Neighbor-Joining dendrogram of *actin* genes (CDS) from *P. patens* and *Actin 7* from *A. thaliana*. The evolutionary distances were computed using the Kimura 2-parameter method. Branch support was provided by bootstrap resampling (10000 replicates). As outgroup an actin related protein (ARP) was included. The dendrogram was built using MEGA5. *Act 7b* Pp3C3_33440V3.1 was called *Act 3* by Weise *et al*., (2006).

**Supplementary Figure S7.**
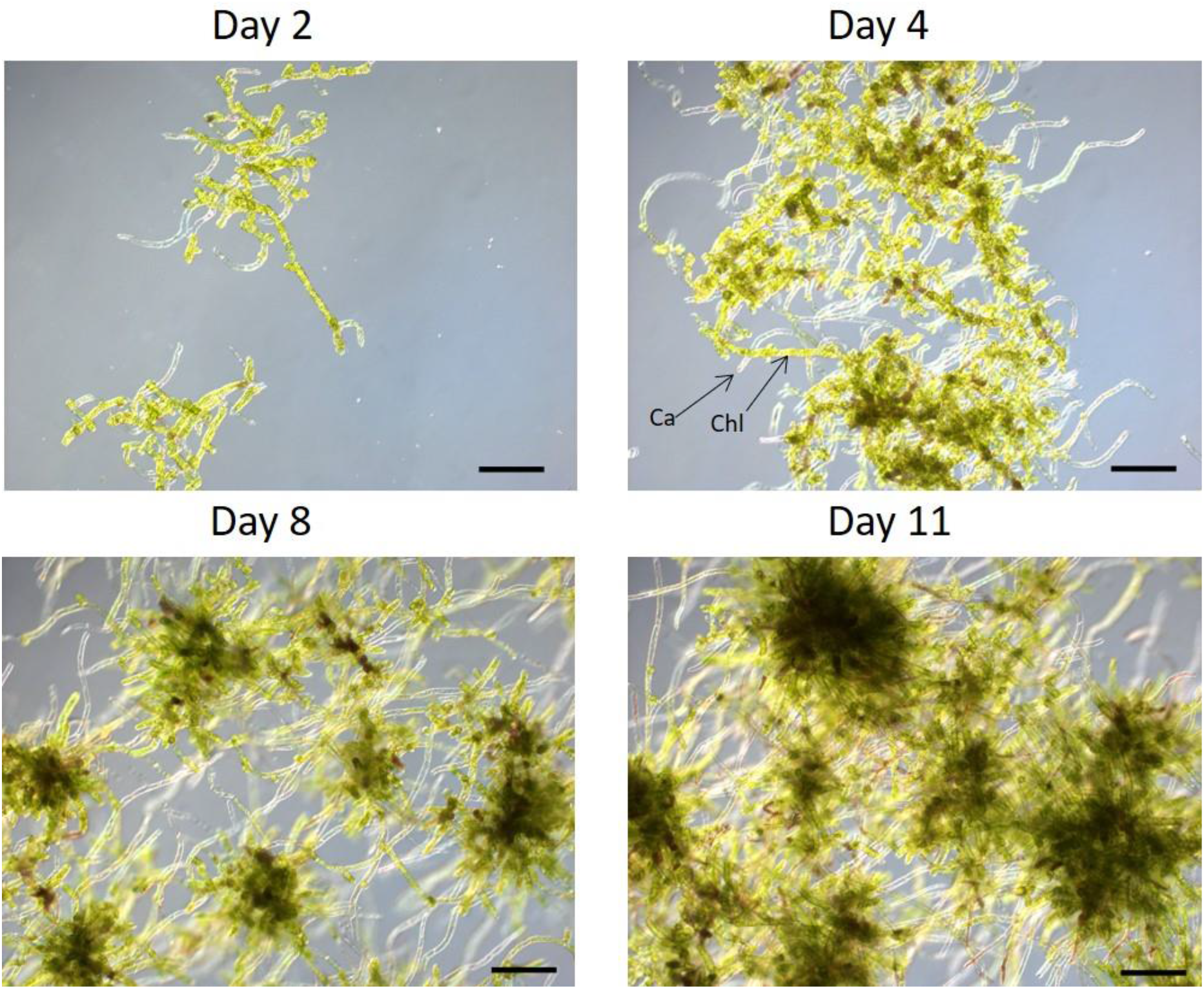
Representative light-microscopic images of Physcomitrella protonema cultivated in 5L stirred-tank bioreactor in batch operation mode with NAA (10 µM) supplementation at day 3. Scale bars: 200 µm. Chloronema predominates the first 2 days, tissue differentiation to caulonema occurs and pellets are visible after 9 days approximately. Chloronema (Chl) and caulonema (Ca) are marked by arrows at day 4.

### Supplementary methods

**Light distribution in the bioreactor and specific growth rate (light as substrate). Methodology described by Evers (1991) (Supplementary eq 1-5)** l_0_ is the incident light [µmol/(m^2^s)], s_x_ is the cell absorption coefficient [m^2^/g] and r_R_ is the cylinder radius [m]. The path length of light (p) is function of a distance from vessel surface (L) and angle of light path (Θ). I(L,c_x_) is the mean light intensity as a function of biomass concentration (c_x_) and location in the vessel.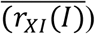 is the overall biomass growth rate considering the light as the limiting substrate.

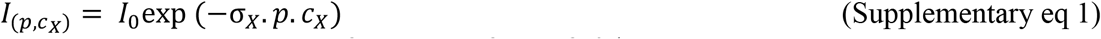

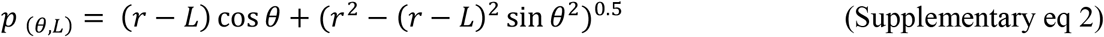

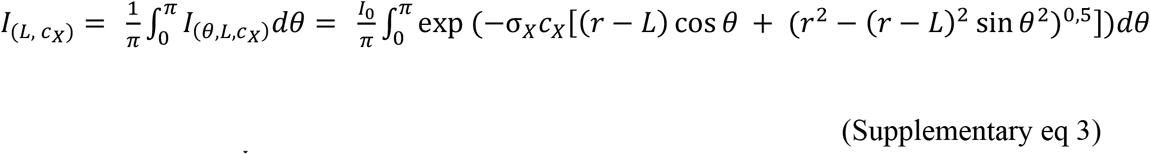

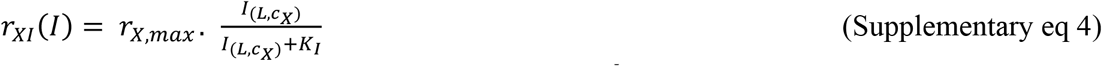

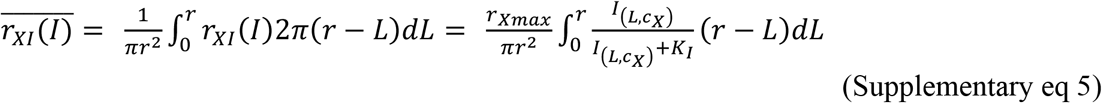

